# Responses of inferior colliculus neurons to notched noises in awake mice: putative neural correlates of auditory enhancement and Zwicker tone

**DOI:** 10.1101/2025.07.09.663932

**Authors:** C-J Hsiao, AV Galazyuk, AJ Norena

## Abstract

Sensory systems are well adapted to constantly changing statistics of the environment and to process specific spectral features of sounds, such as spectral notch (i.e. low energy frequency band) embedded in broadband stimuli. Spectral notches can be added to the stimulus spectrum due to filtering by the outer ear, and can be used as monaural cues related to head or pinna position for localizing sound sources. In addition, broadband sounds with spectral notch are known to produce auditory enhancement, a perceptual phenomenon in which a target within a spectrally notched masker can become salient if preceded by a copy of the masker. Notched noise can also produce an auditory illusion, called Zwicker Tone (ZT), which is perceived immediately after stimulation and whose pitch corresponds to the spectral notch. The present study aimed to further investigate the mechanisms of auditory enhancement, including those of ZT, in the inferior colliculus of awake mice. We show that neural activity can be strongly suppressed during NN stimulation and enhanced immediately after NN stimulation. These effects depend on notch center frequency relative to the best frequency of neurons, stimulus level and notch width. Our results are consistent with the mechanisms described for post-inhibitory rebound in the central auditory system: NN could hyperpolarize the membrane potential, which can then activate several cationic conductances, leading to a rebound of neural activity. We discuss auditory enhancement and ZT as collateral effects of an essential neural mechanism aimed at enhancing the central representation of acoustic spectral contrasts.

## Introduction

Sensory systems are well adapted to constant changes in the statistical characteristics of the environment. For example, sensory information is processed in such a way as to preserve perceptual stability despite significant variations in spectro-temporal energy in the environment (Stilp and Assgari, 2018). The ability of sensory systems to compute the statistics of the sensory context can also play a role in emphasizing the processing of novel events (Grimm et al., 2016). Furthermore, neural processing is optimized to maximize neural resources and coding efficiency in a given sensory context. In this context, it has been suggested that the rate-level function of neurons (i.e., neuronal sensitivity to sound level) can adapt quickly (over seconds or minutes) to the distribution of stimulus levels : the slope of the rate-level function tends to be maximal (maximum coding accuracy) around the level corresponding to the peak distribution (Dean et al., 2005). Intuitively, these changes can be understood as follows: the system allocates greater coding resolution where a given dimension of the stimulus is more prevalent (Noreña, 2011; Rieke and Rudd, 2009).

It is also well known that perception, including sensory illusions, depends heavily on context and recent stimulation history (Yasoda-Mohan et al., 2025; Zhang et al., 2025). In the auditory modality, the phenomenon known as “auditory enhancement” illustrates this characteristic of the auditory system, which consists of processing sensory information according to context (Byrne et al., 2011; Viemeister and Bacon, 1982; Wang and Oxenham, 2016). Classically, the paradigm is defined as follows: a target within a spectrally notched masker (complex stimulus) can « pop out » or become salient if preceded by the masker (without the target). Both target and masker start and end at the same time, the target frequency corresponds to the notch center frequency and the masker plus target stimulus is presented quickly (within a few hundreds of msec) after the masker sound. In short, the spectral region where there was no energy present is specifically enhanced immediately after stimulus presentation. This paradigm has been shown to affect speech perception (Stilp, 2019; Stilp and Assgari, 2018). The mechanisms of this auditory enhancement are largely unknown and speculative. First, it was suggested that the increase in salience of the target could result from neural adaptation of the spectral region stimulated by the masker (Stilp and Assgari, 2018; Summerfield et al., 1987). According to this hypothesis, the neural representation (i.e., absolute value of the firing rate) of the target is not increased, but is relatively enhanced compared to the activity corresponding to the surrounding frequency regions (masker), where neural activity is supposed to be reduced by neural adaptation. However, this explanation does not account for all results collected in psychoacoustic studies. In particular, an « enhanced » target can produce more forward masking than a control target on a tone detection task, suggesting that target enhancement is absolute (true gain) and not relative. Another hypothesis, known as adaptation of lateral inhibition, has been then advanced (Byrne et al., 2011; Carcagno et al., 2012; Nelson and Young, 2010; Viemeister and Bacon, 1982; Wang and Oxenham, 2016). Lateral inhibition, which is directed towards the spectral notch of the masker from the frequency regions surrounding the target, is supposed to be adapted by the presentation of the masker alone. Then, when the target and the masker are presented quickly after the masker alone, the target frequency undergoes less inhibition and is therefore enhanced compared to the same stimulus not preceded by the masker alone. Neural correlates (amplitude increase of steady-state responses) of auditory enhancement have been reported in human subjects, suggesting that the spectral region within the spectral notch is enhanced (Mehta et al., 2021).

While conducting some psychoacoustic experiments, Zwicker reported hearing a “negative afterimage”, later called the « Zwicker tone » (ZT), after being presented with notched noise (NN - broadband noise with a spectral notch) (Zwicker, 1964). He described the ZT as a short « bing », with a pitch located within the spectral gap, and lasting a few hundreds of msec to seconds after the presentation of NN for a few seconds to 1 minute, respectively. The NN is also followed by auditory enhancement, namely a decrease of absolute thresholds close to 10 dB within the spectral notch region (Norena et al., 2000; Wiegrebe et al., 1996). The ZT produced after NN presentation suggests that the mechanisms underlying auditory enhancement could also enhance the background (spontaneous) neural activity (Norena et al., 2000), in addition to enhancing the stimulus-induced activity. In other words, the ZT could result from a transient increase of neural activity, i.e. a specific type of offset responses, immediately after NN presentation. It has been suggested that ZT could be a form of transient tinnitus, or equivalently, that NN could create the conditions (i.e., central changes) that could ultimately lead to transient tinnitus (Norena et al., 2000; Parra and Pearlmutter, 2007; Schilling et al., 2023). Recently, we investigated the cortical responses in anesthetized and awake guinea pigs during and after notched noise stimulation. Interestingly, more offset responses were observed after NN stimulation compared to the control condition (white noise presentation). However, offset responses were usually relatively short (<100 msec) and therefore were unlikely to account for the ZT sensation (which can last for a few hundreds of msec to sec) (Schilling et al., 2023).

The present study aimed to further investigate the mechanisms of auditory enhancement, including those of ZT, in the inferior colliculus (IC) of awake mice. We chose to focus on the IC for several reasons. First, the IC is a very important relay in the central auditory system, where many cell types and neural responses have been described, and where inhibitory processes are also very strong (Gittelman et al., 2012; LeBeau et al., 2001). Moreover, most neurons in the IC present spontaneous activity in silence and tonic discharges during acoustic stimulation (Voytenko and Galazyuk, 2010). Hence, this pattern of discharges allows us to investigate the net inhibitory or excitatory effect of a given stimulus (white noise and NN in our study) on neuronal activity during and after acoustic stimulation. This is important regarding the hypothesis suggesting that inhibition (i.e. adaptation of inhibition) is supposed to play a key role in auditory enhancement and potentially ZT generation (Byrne et al., 2011; Viemeister and Bacon, 1982; Wang and Oxenham, 2016). Also, NN and other stimuli with a missing spectral region are well designed to modify the balance between excitation and inhibition. Indeed, many neurons in IC have lateral inhibitory side bands adjacent to their excitatory tuning curve (Xie et al., 2008). Notch noises with notch center corresponding to the best frequency of neurons are well suited to recruit inhibitory side bands, while not stimulating the central excitatory region. As a consequence, the net effect of NN may be inhibitory and neurons may present low activity, if any, during NN presentation. Auditory enhancement (and ZT) may then reflect a post-inhibitory rebound (Kopp-Scheinpflug et al., 2018) after NN presentation increasing both spontaneous and stimulus-evoked neural activity within the frequency region of the notch. Hence, the present study is aimed at investigating the net effect between excitation and inhibition, and potential non-linearities (including post-inhibitory rebound) during and after NN presentation. The effects of various characteristics of NN have been studied, namely the notch center frequency (relative to the neurons’ best frequency), notch width, and intensity levels. We show that neural activity can be strongly suppressed during NN stimulation and enhanced immediately after NN stimulation. These effects were maximal at stimulus level of 60 dB SPL, notch width of 1 octave and when the notch center frequency corresponded to neuron’s best frequency. Our results are consistent with the mechanisms described for post-inhibitory rebound in the central auditory system (Kopp-Scheinpflug et al., 2018, 2011). We interpret auditory enhancement and ZT as a collateral effects of an essential neural mechanisms aimed at enhancing spectral contrasts within broadband sounds.

## Material and methods

### Subjects

A total of 9 CBA/CAJ mice (4 male and 5 female), aged between 6 and 14 months, were used in this study. Animals were housed in pairs in a colony room maintained at 25°C with a 12-hour light/dark cycle. Animal procedures were approved by the Institutional Animal Care and Use Committee (IACUC) at Northeast Ohio Medical University.

### Surgery

Each mouse was anesthetized during surgery with 1.5–2.0% isoflurane. A midline scalp incision was made and the tissue overlying the cranium was removed. A small metal rod was then affixed to the skull using dental cement (C&B Metabond, Japan). After a recovery period of at least two days, animals were habituated to a custom holding device inside a single-walled sound-attenuating chamber. The holding device consisted of a custom-made plastic tube and a small metal holder. During electrophysiological recordings, the animals’ ears remained unobstructed to allow for free-field acoustic stimulation.

### Acoustic stimulation

The stimulation protocol consisted of three components. First, spontaneous firing rate (SFR) was measured during a 10-second silence recording window without sound stimulation. Second, a rapid sequence was used to estimate the best frequency (BF, frequency at which firing rate is highest) consisting of presenting tones (100-130 msec duration) across a broad frequency range (2-64 kHz in 1/4-octave steps; 21 total steps) presented at 60 dB SPL and at least twice. Third, a sequence of 1-second white noise or notched noises centered at the neuron’s BF and non-BF followed by 1 sec silence were presented at least 15 times. White noise (WN) and notched noises (NN) were generated using Matlab© custom program at a sampling rate of 195312 Hz. Notched noises were obtained from FFT filter. First, a WN was converted in the frequency domain. Filtering was applied in the frequency domain by zeroing the amplitude in the frequency band corresponding to the notch. Finally, the signal is transformed into the time domain by using an inverse FFT. The noise stimuli had a linear rise and fall ramp of 10 msec.

All acoustic stimuli were generated by a Tucker-Davis Technologies system 3, which included an RX6 multifunction processor, a PA5 programmable attenuator and SigGenRP software. The signals were amplified using an HCA-750A amplifier (PARASOUND) and delivered via a free-field loudspeaker (LCYK100 Ribbon Tweeter, Madisound) positioned 10 cm in front of the animal. Stimuli were calibrated using a 1/4-inch free-field microphone (Type 4939, Brüel and Kjær) placed at 10 cm in front of the speaker with a conditioning amplifier (NEXUS 2690-A, Brüel and Kjær) sampling at 195.3 kHz to ensure accurate sound pressure levels measurements across frequencies.

### Extracellular Recordings

Recordings were obtained from both the right and left inferior colliculus (IC) in unanesthetized mice housed in a single-walled sound-attenuating chamber (Industrial Acoustics Company, Inc.). Throughout each recording session (3–4 hours), animals were periodically offered water and continuously monitored for signs of discomfort. After the end of each recording session, the exposed skull was sealed with a Kwik-Sil silicone elastomer plug (World Precision Instruments) and the animal was returned to its home cage. All experiments were conducted on adult mice at least two months of age. Recordings were performed every other day for up to two weeks, after which the animal was euthanized via intraperitoneal injection of Fatal-Plus. No sedative drugs were used during recording sessions. If any signs of discomfort were observed, the recording session was immediately terminated, and the mouse was returned to its cage.

Recording electrodes were inserted through a small craniotomy overlying the IC. Extracellular single-unit recordings were made using quartz glass micropipettes (10–20 MΩ impedance, 2–3 μm tip) filled with 0.5 M sodium chloride. Electrodes were fabricated with a P-2000 horizontal micropipette puller (Sutter Instrument) and positioned using a precision digital micromanipulator (MP-225, Sutter Instrument) under a surgical microscope (Leica MZ9.5). Electrode position was tracked using digital micrometer readouts reference to a fixed point on the brain surface.

Recordings were restricted to the central nucleus of the IC based on the electrode depth. Electrode advancement was controlled remotely via a piezoelectric microdrive (Model 660, KOPF Instr.) located outside the sound-attenuating chamber. Action potential was amplified using a preamplifier (Model 2400A. Dagan), monitored audibly with an audio monitor (AM7, Grass Instruments) and visually with a digital oscilloscope (DL3024, YOKOGAWA), digitized and stored on a computer via an EPC-10 digital interface and PULSE software (HEKA Elektronik) at a 100 kHz sampling rate.

### Statistics

Neuronal response patterns can vary considerably, suggesting that cellular activity should not be averaged across a large number of cells, as this could cause important information to be lost. On the other hand, it is also more convincing to use statistics to evaluate differences between various conditions at the group level. Hence, we chose a mixed way of reporting results : individual results are reported so the diversity of neural patterns are well illustrated. Firing rate was obtained using a sliding gaussian window of 20 msec (standard deviation=2.5).

On each of the individual results shown in the figures, a horizontal gray rectangles was drawn illustrating the values of the average spontaneous activity (calculated from the time window between 400 and 650 ms after stimulus presentation) plus or minus twice the standard deviation. This simple statistical approach makes it easy to compare the neural activity of a given cell to its own spontaneous activity, which is also easy to illustrate.

We also compared NN and WN conditions using a statistical approach we already used and described earlier (Parameshwarappa et al., 2024). We used a statistical test that is nonparametric (independent from data distribution) and straightforward to deal with the multiple comparisons problem (here comparisons over time, at each time sample – every msec) (Maris and Oostenveld, 2007). The approach is first aimed at calculating the threshold for statistically significant clusters. It was done in the following way :

1. The data from NN and WN conditions were grouped in a single set.
2. We then generated a random partition for condition 1 (WN) and condition 2 (NN) from the set built in step 1.
3. The test statistic (t-test) was calculated on this random partition
4. All values below -2 and above 2 were considered as crossing the threshold of “significance”. The values above (or below) threshold can form clusters (adjacent time samples where the test statistics cross the threshold). The t-test values within each cluster were then summed and the largest and smallest were kept.
5. Steps 2 to 4 are repeated 1000 times.
6. The threshold for the t-test was then derived from the distribution of largest (and smallest) values of the cluster-level statistics: 2.5th and 97.5 quantile.

The different NN conditions were compared to the corresponding control condition using the t-test. Then, the clusters (connected samples crossing the threshold) were considered significant if the sum within a cluster crossed the threshold obtained in the procedure described above.

## Results

Overall, we could record from 99 cells in the IC of awake mice. Onset responses were observed in 43 cells (for at least one stimulus condition). Offset responses were detected in 16 cells after WN and 55 cells after NN (for at least one stimulus condition). Onset and offset responses were both present on a same recording in only 18 cells.

### Individual examples

In our set-up, we could record from a given cell for a few minutes and rarely for more than 15 mins consecutively. That is why we did not test all stimulus conditions in all cells.

Figure 1 shows the frequency response maps (see methods) obtained for the nine cells whose responses evoked by WN and NN are shown in Figures 2-10. Most cells present relatively narrow excitatory responses over frequency surrounded by lateral inhibitory side bands.

**Figure 1.**
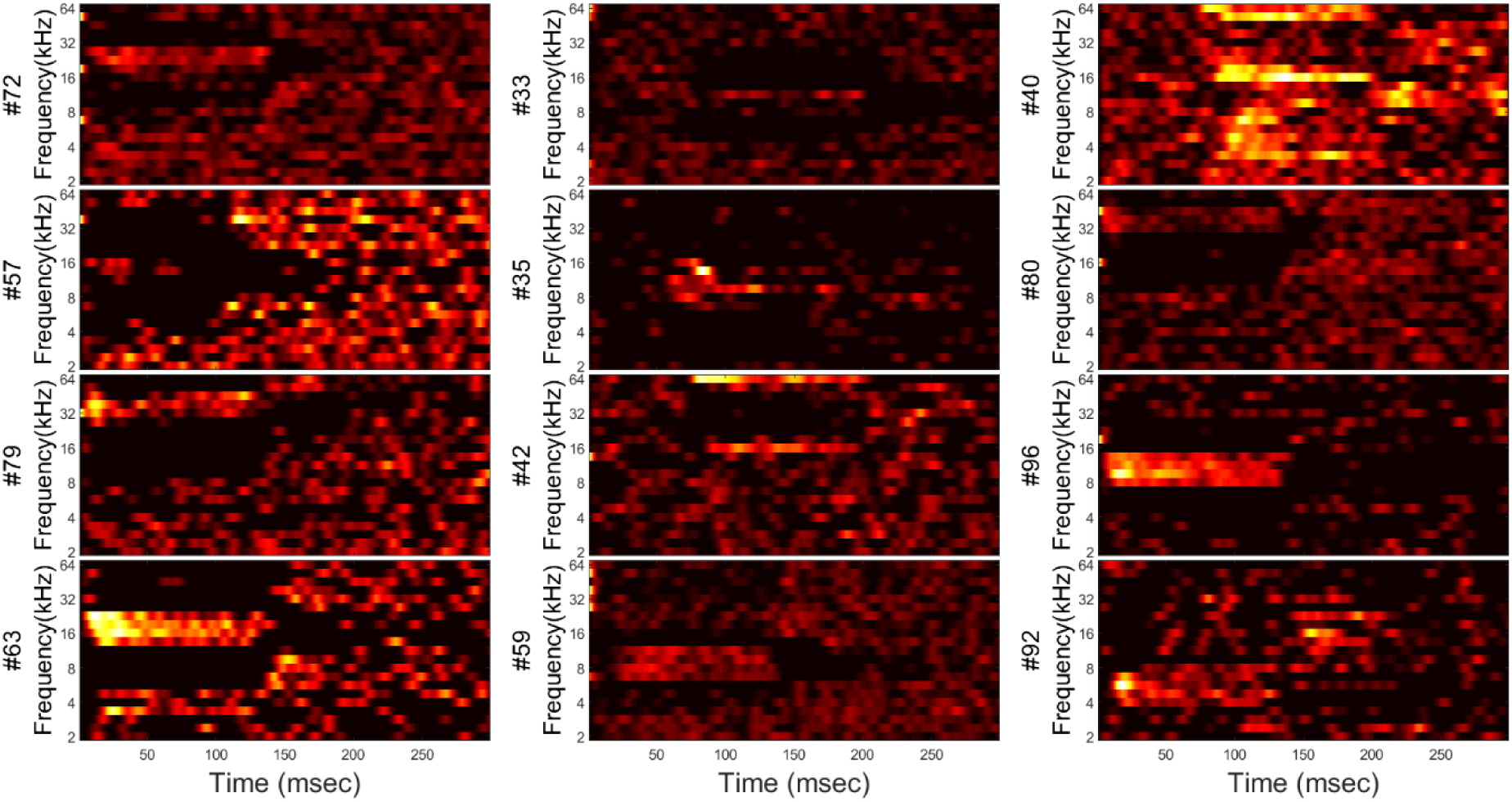
Frequency response as a function of time for all cells whose responses to NN and WN are shown in Figures 2-10. Tone pips (100-130 ms, frequency from 2kHz to 64kHz, step of ¼ octave) were presented at 60 dB SPL. In a few examples, tone pips were preceded by 50-70 ms silence. Cell numbers are shown on the left side of each frequency response. The most common pattern of responses is narrow excitatory area surrounded by lateral inhibitory sidebands (#72, #57, #79, #63, #33, #35, #59, #80, #96).

**Figure 2.**
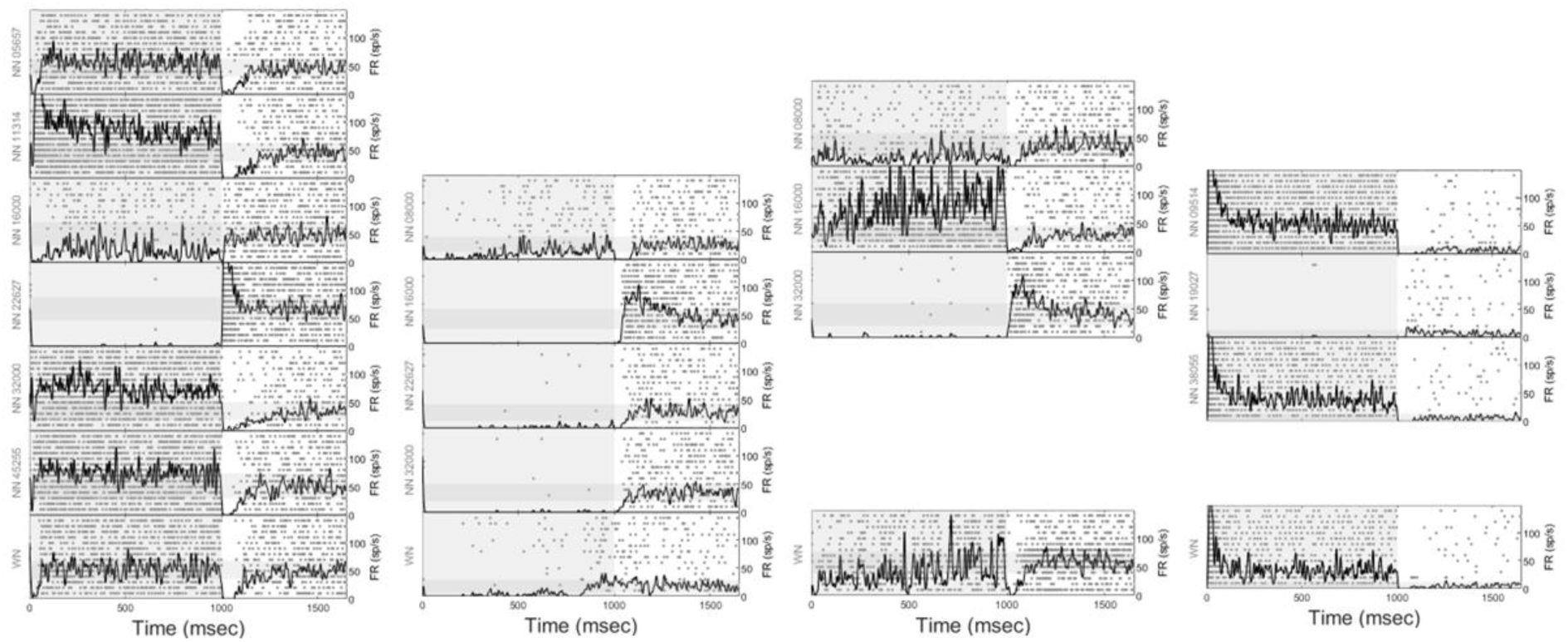
From left to right, the figure shows the pattern of responses for 4 cells (#72-BF=22627Hz, #57-BF=16000Hz, #79-BF=32000Hz, #63-BF=19027Hz). The notch center frequency (for NN, WN otherwise) is indicated on the left side of each figure. Each gray dot corresponds to an action potential (each row corresponds to a sound sequence, 15 repetitions are shown). The black line corresponds to the firing rate (right y-axis) calculated using a sliding gaussian window of 20 msec. The light gray area corresponds to the stimulus presentation, and the darker gray horizontal band (from 0 to 1650 msec) to the mean spontaneous activity +/- 2 * standard deviation (derived from the time window between 400 and 650 msec after sound stimulation). Missing panels are absent when we did not have time to collect the corresponding data.

Figure 2 shows four individual examples illustrating different pattern of responses during and after noise stimulation presented at 60 dB SPL. The first example (first column on the left, BF=22627 Hz) shows a typical pattern of neural responses during and after WN stimulation for different conditions of NN stimulation (various notch center frequencies). This cell has a spontaneous firing rate at around 50 spikes / sec, and a tonic response slightly above this value. The cell does not show any onset responses to the sound, except for the NN condition with a notch center frequency at 11314 Hz (NN_11314Hz). Interestingly, the cell presents a tonic response during all stimulus conditions, except for the NN_22627Hz (BF): during the stimulation, neural activity is almost completely suppressed. Moreover, immediately after the stimulation (NN_22627Hz), the offset response is large and relatively long lasting (around 200 msec). On the contrary, neural activity for most other stimulus conditions are suppressed immediately after stimulus presentation. One notes the high frequency specificity of this neural activity pattern : for NN stimulations with notch center frequency at half-an-octave above or below the BF, the firing is tonic during NN stimulation and there is no offset response. It is interesting to note that the discharge pattern of NN_16000Hz lies halfway between that of NN_22627 (BF) and that of others NN, whose notch center lies at other frequencies: a certain sustained discharge rate is observed during stimulation and no suppression of the discharge rate immediately after stimulus presentation. Examples shown in second (BF=16kHz) and third column (BF=32kHz) illustrate other pattern of responses. Sustained activity is dramatically reduced during NN stimulation. The example in second column illustrates the sensitivity of offset responses to neural suppression during stimulation: offset responses are observed only when the sustained activity is abolished (no spikes at all). The example in third column illustrates the diversity of sustained activity: sustained activity is maximal for NN_16000Hz. The last example (fourth column) shows strong suppression for NN_19027Hz (BF) but no offset responses.

Figure 3 shows another example of cell during and after WN and NN stimulation as a function of intensity level (BF=8000 Hz). At 40 dB SPL, the pattern of neural activity is similar across the stimulus conditions (buildup pattern of activity), except for NN_8000Hz (BF) where a small offset response is visible. At 60 and 80 dB SPL, the neural activity during stimulation with NN_8000Hz is almost completely suppressed, and an offset response is present (larger at 60 dB SPL compared to 80 dB SPL). The pattern of offset responses at 60 dB SPL and NN_8000Hz is interesting as it presents two phases: shortly after the stimulation, a sharp offset response is visible, and after a brief decrease in the discharge rate, a more spread-out offset response can be observed. This result suggests that offset responses result from different mechanisms (different cationic conductances, see discussion) (Kopp-Scheinpflug et al., 2018). One also notes that onset responses vary with stimulus conditions (spectral shape and intensity level).

**Figure 3.**
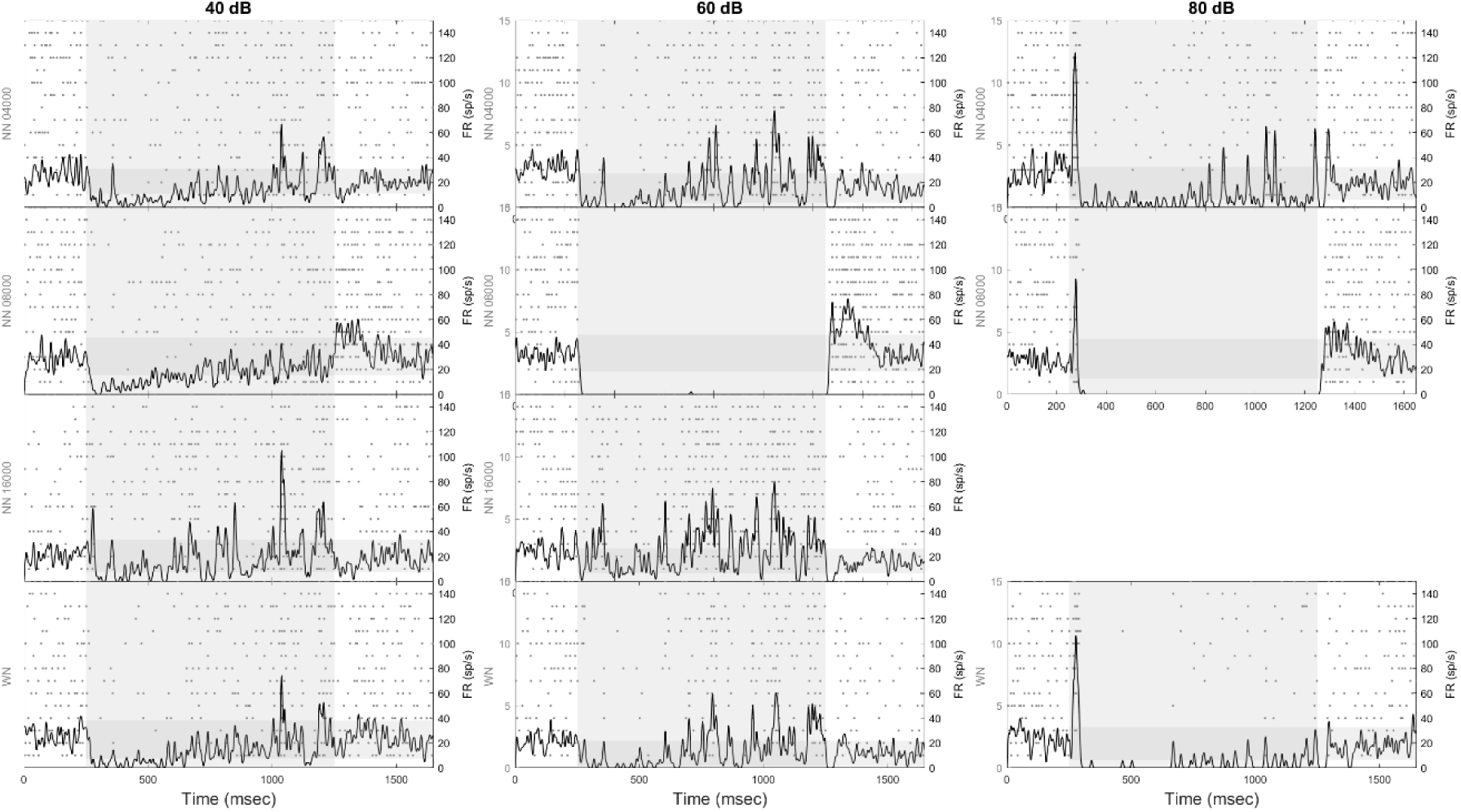
The figure shows the pattern of responses for cell #33. Same than Figure 2 otherwise.

Figure 4 shows another example (BF=8000Hz), comparable to the previous one, illustrating different patterns of onset and offset responses. First, offset responses were observed at NN_8000Hz (BF) but also at NN_6727Hz. Second, while the offset responses spread over more than 100 msec at 40 dB SPL, it is sharper after NN_8000Hz presented at 60 dB SPL, and are small at 80 dB SPL. In this example, we note that onset and offset responses are both present for a few conditions, especially NN_6027Hz presented at 60 dB.

**Figure 4.**
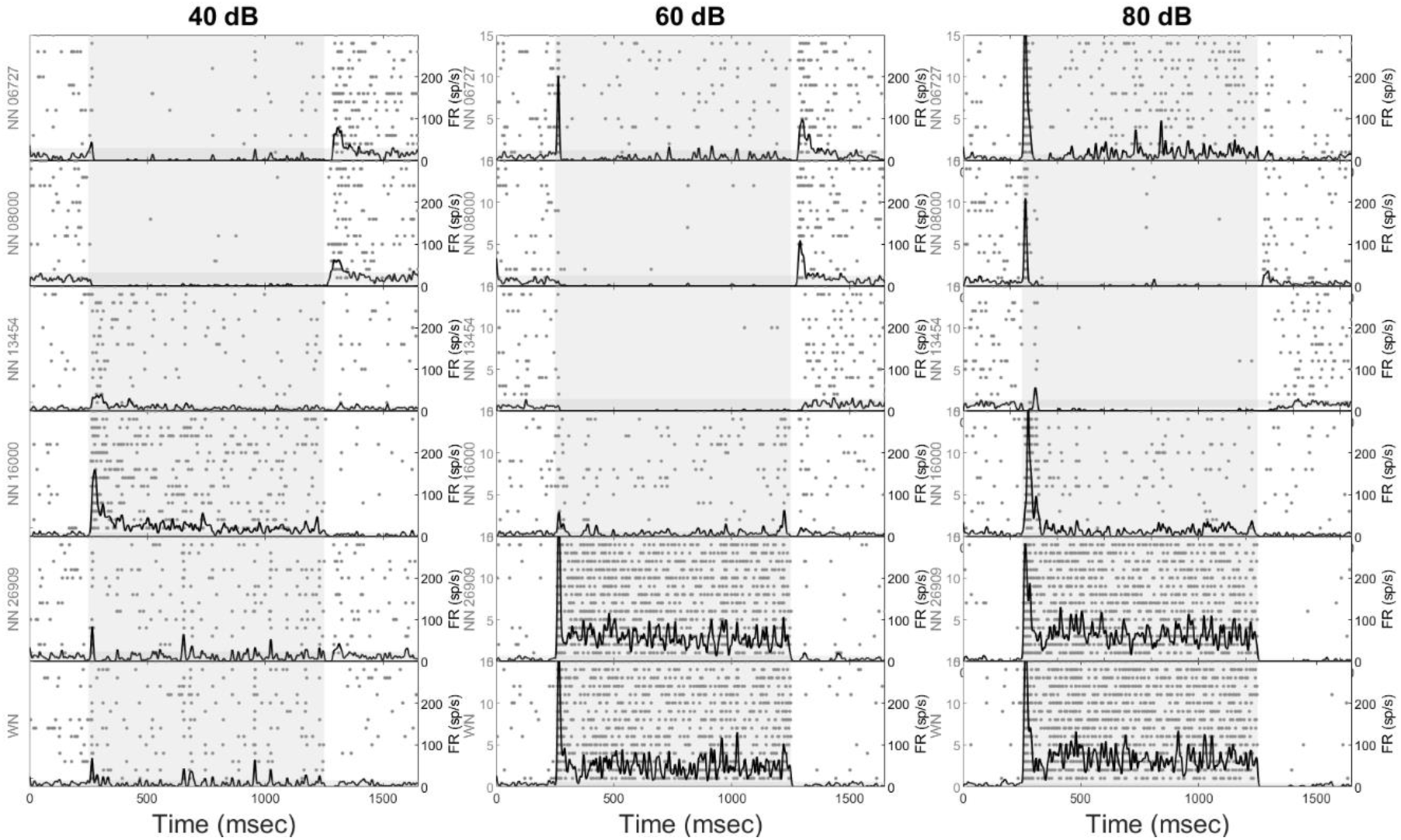
The figure shows the pattern of responses for cell #35. Same than Figure 2 otherwise.

The example shown in Figure 5 has been chosen as it illustrates the changes in the pattern of sustained activity with NN and NN. For WN and NN_16000Hz, the firing exhibit buildup pattern at 40 dB SPL, sustained for NN_32000Hz, while activity is strongly suppressed for NN_64000Hz (BF). Onset and offset responses also vary as a function of notch center frequency and level. Interestingly, while there was no onset response but a small offset response for NN_64000Hz at 60 dB SPL, a small onset response is present but there is no offset response at 80 dB SPL.

**Figure 5.**
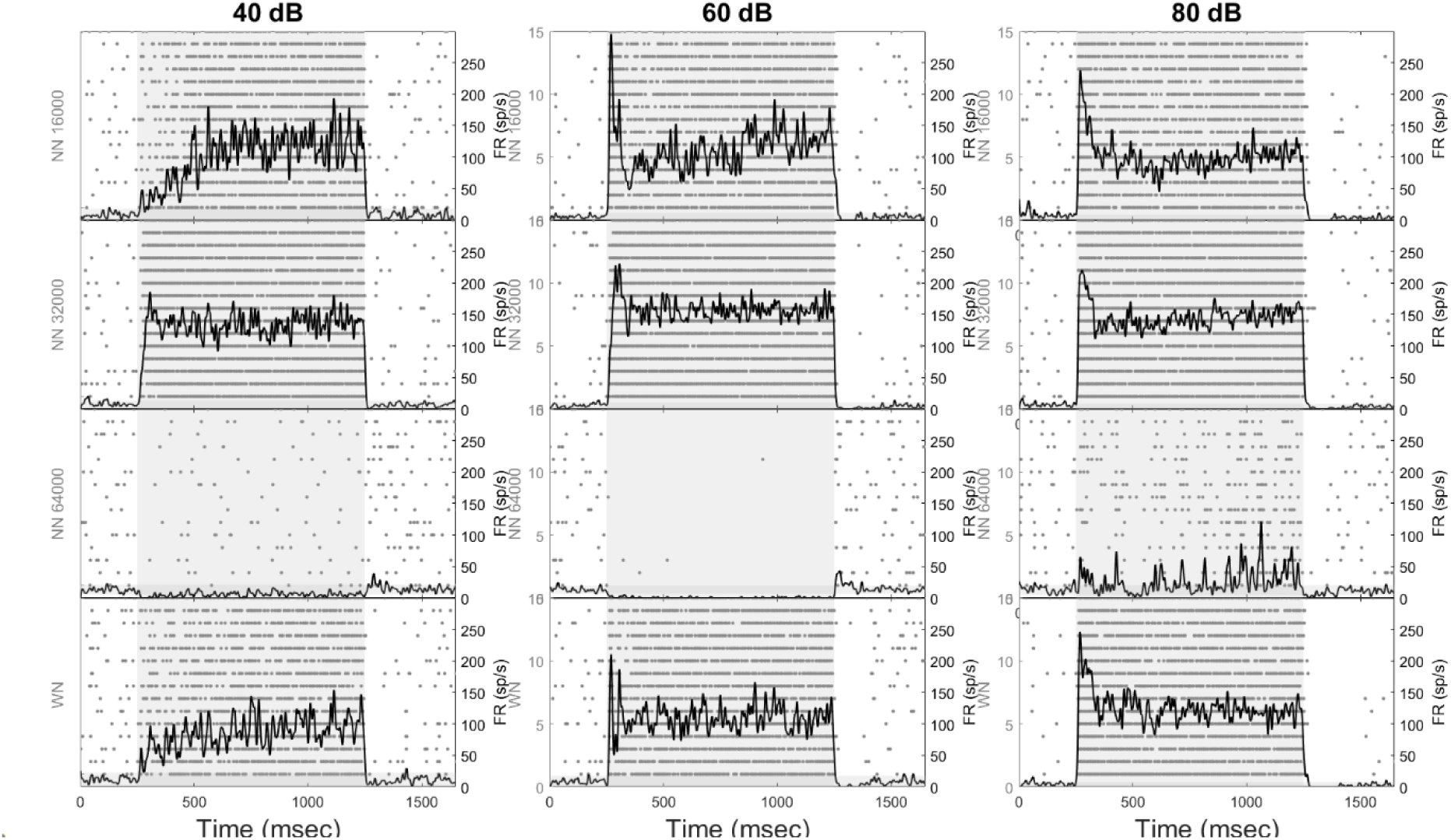
The figure shows the pattern of responses for cell #42. Same than Figure 2 otherwise.

The cell shown in Figure 6 illustrates another pattern of firing as a function of stimulus conditions. First, the firing rate is higher for NN_8000Hz (BF) compared to WN at 40 dB SPL. At 60 dB SPL the pattern of firing is the other way around: firing rate is suppressed during NN_8000Hz and enhanced during WN. Moreover, this example shows a large and prolonged suppression of activity immediately after stimulation, except after NN_8000Hz (BF) where the offset response is large (at 60 dB SPL). At 80 dB SPL, the pattern of offset responses is also interesting: there is a sharp offset response (although below the background activity), followed by a decrease of activity and a slow return to the (high) background activity. This cell (see also Figure 2) illustrates what can be an important property of the auditory system: the frequency band where energy was missing in the recent stimulation history is enhanced, while, and on the contrary, the frequency region where energy was present is inhibited. The suppression of neural activity after stimulation can last up to 200 msec, which is important and functionally potentially meaningful (see discussion).

**Figure 6.**
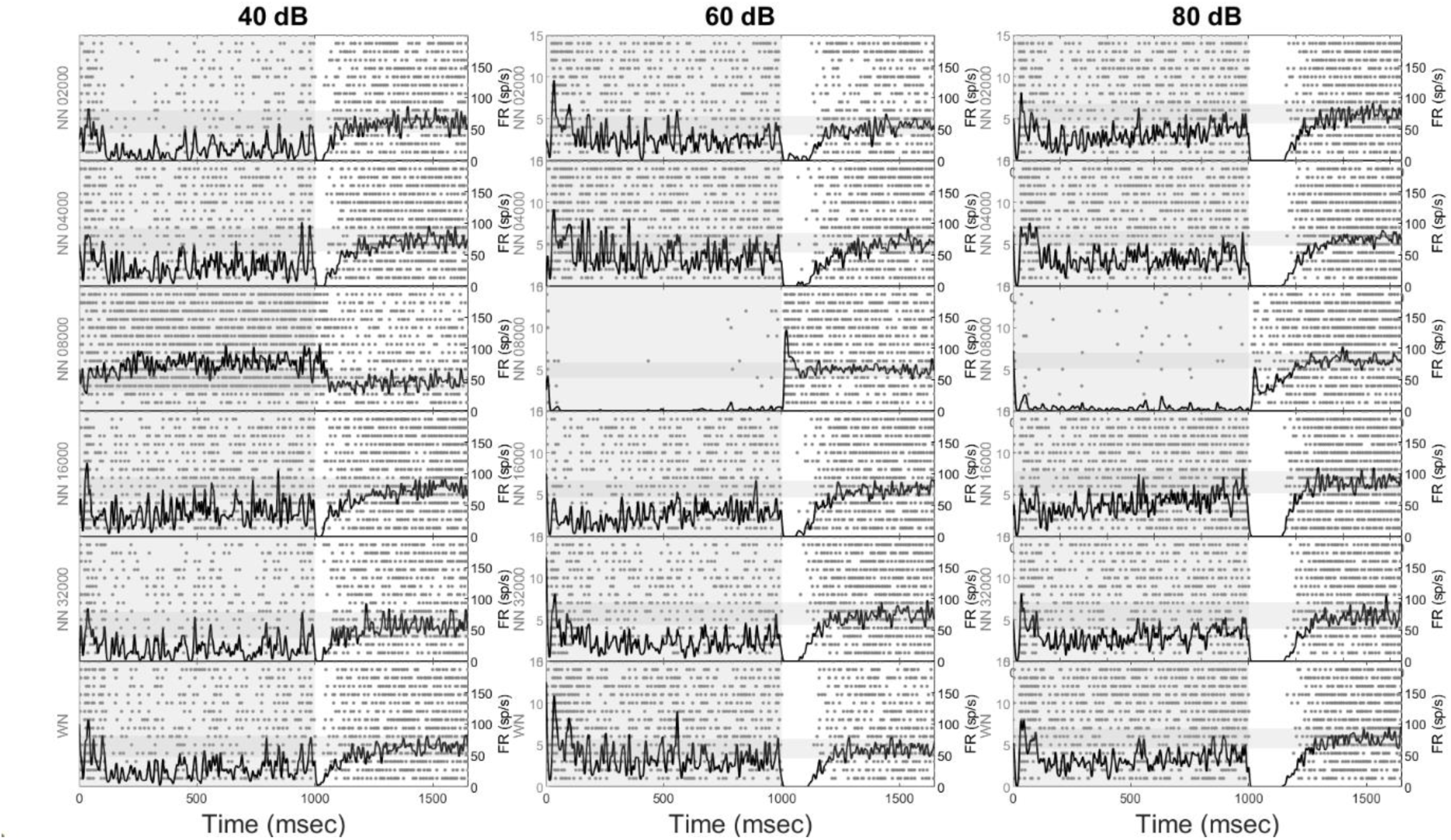
The figure shows the pattern of responses for cell #59. Same than Figure 2 otherwise.

Figure 7 shows an example of cell where onset and offset responses were both present for the same stimulus conditions. Onset response was present in all conditions, while offset response was present at BF (NN_64000Hz) and half-an-octave below BF (NN_45255Hz). Offset responses are observed while neural activity is present during NN stimulation. However, neural activity in the last 500 msec of stimulation is at (NN_45255Hz) or below (NN_64000Hz) the spontaneous activity.

**Figure 7.**
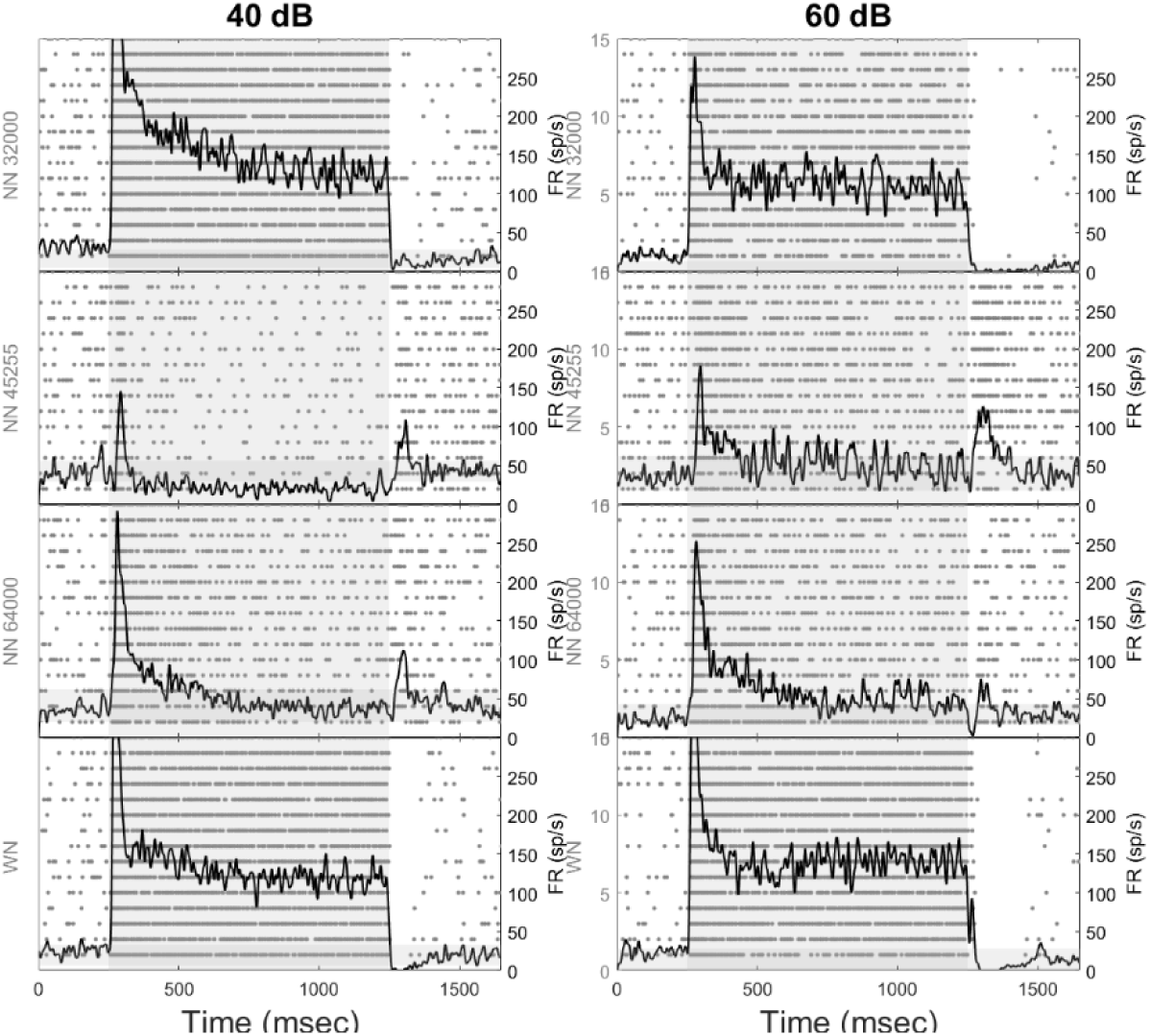
The figure shows the pattern of responses for cell #40. Same than Figure 2 otherwise.

Figures 8-10 show examples of cells where the effects of varying the notch width have been studied (stimulus level was fixed at 60 dB SPL). Overall, the pattern of neural activity is consistent with the examples shown above: neural activity is reduced during NN presentation and enhanced after NN presentation. Example shown in Figure 8 (BF=32000Hz) shows that the notch width of one octave is optimal for suppressing and increasing neural activity during and after NN_32000Hz presentation, respectively. We want to emphasize again in this example, the sensitivity of neural responses to stimulus conditions: both neural suppression and offset response during and after NN presentation are obtained for NN_32000Hz, but not for NN_22627Hz and NN_45255Hz. This example also shows strong suppression of neural activity immediately after stimulus presentation (except for NN_32000Hz). Example in Figure 9 shows another cell with suppression and offset response for NN_8000Hz (BF). Offset responses are observed for NN_8000Hz and NN_16000Hz (for the condition with notch width of 1 octave) but they differ in their pattern of firing: offset is sharp after NN_16000Hz, while it is more spread-out after NN_8000Hz. The last example (Figure 10) illustrates the complexity of neural responses when the notch center frequency and notch width are varied (BF = 4000 Hz). The cell presents both onset and offset responses (NW=0.5 oct, NN_5657Hz and lower), onset responses only (NW=0.5 oct, NN_8000Hz), or offset responses only (NN_4000Hz, NW= 1 octave).

**Figure 8.**
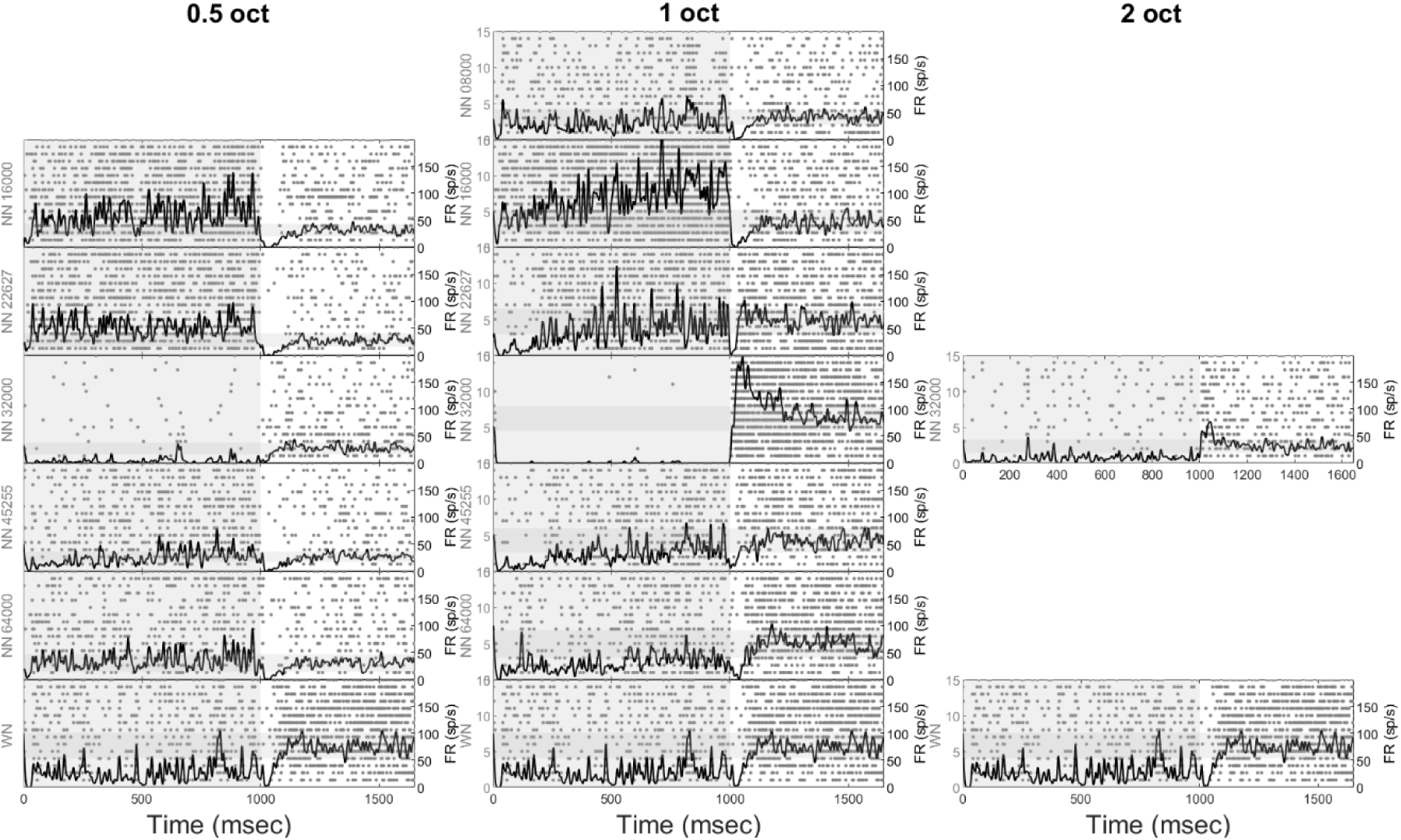
The figure shows the pattern of responses for cell #80. Same than Figure 2 otherwise.

**Figure 9.**
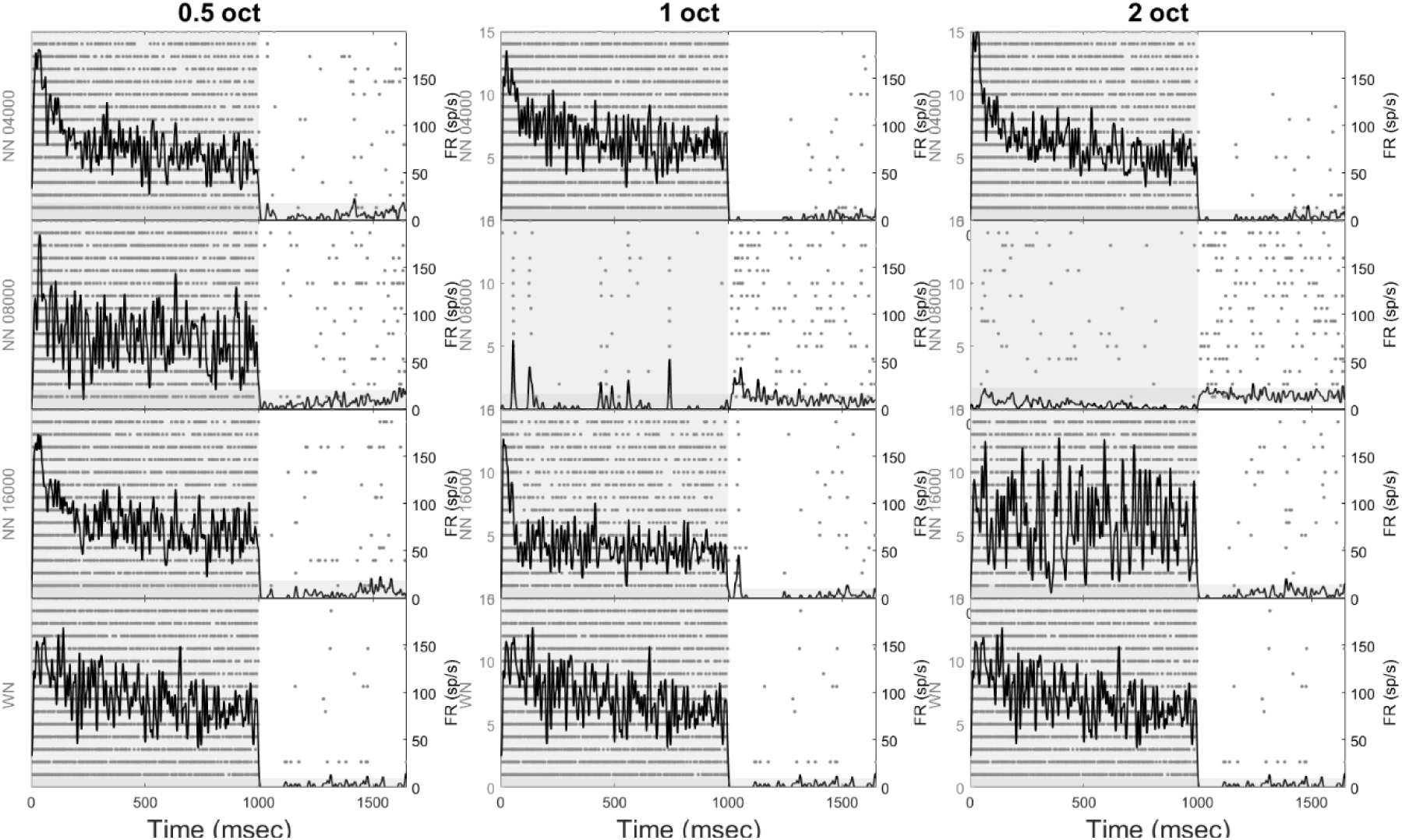
The figure shows the pattern of responses for cell #96. Same than Figure 2 otherwise.

**Figure 10.**
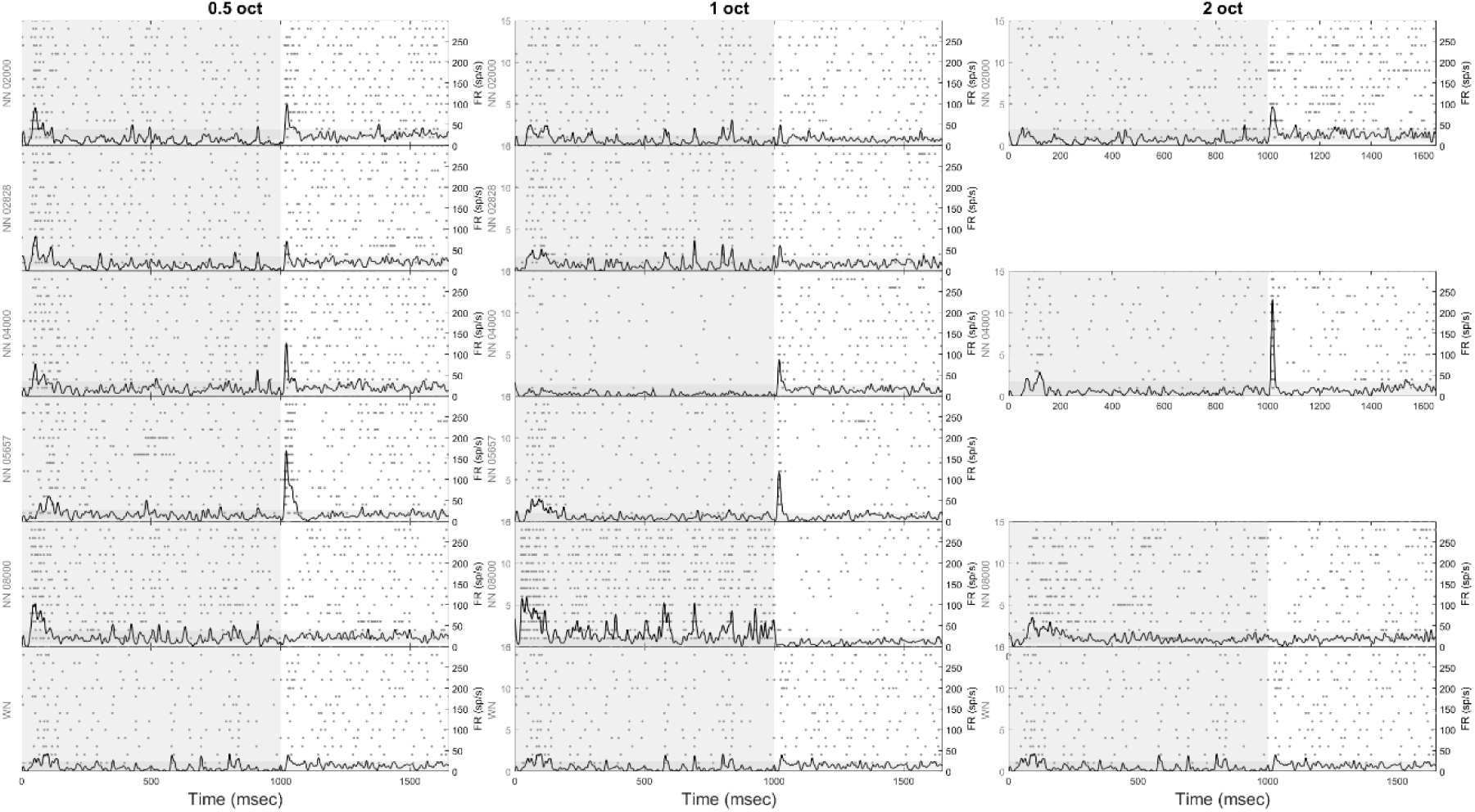
The figure shows the pattern of responses for cell #92. Same than Figure 2 otherwise.

### Group data

Figure 11 shows the averaged data obtained from 39 cells in which we have been able to document the neural activity in response to WN and NN whose center frequency of the notch was located at BF (NN_BF), one octave above (NN_BF+1oct) or below BF (NN_BF-1oct). Each NN condition was compared to WN using the nonparametric test described in the method section. The horizontal black segments indicate the significant clusters (p<0.05). Neural activity was significantly reduced during the NN_CF presentation compared to WN, and the offset response was enhanced after NN_CF compared to WN. The duration of the offset response after NN_CF was spread-out, lasting around 200 msec. In average, there was no offset response for other conditions (WN, NN_BF+1oct and NN_BF-1oct), the activity was even reduced compared to the background activity for almost 200 msec.

**Figure 11.**
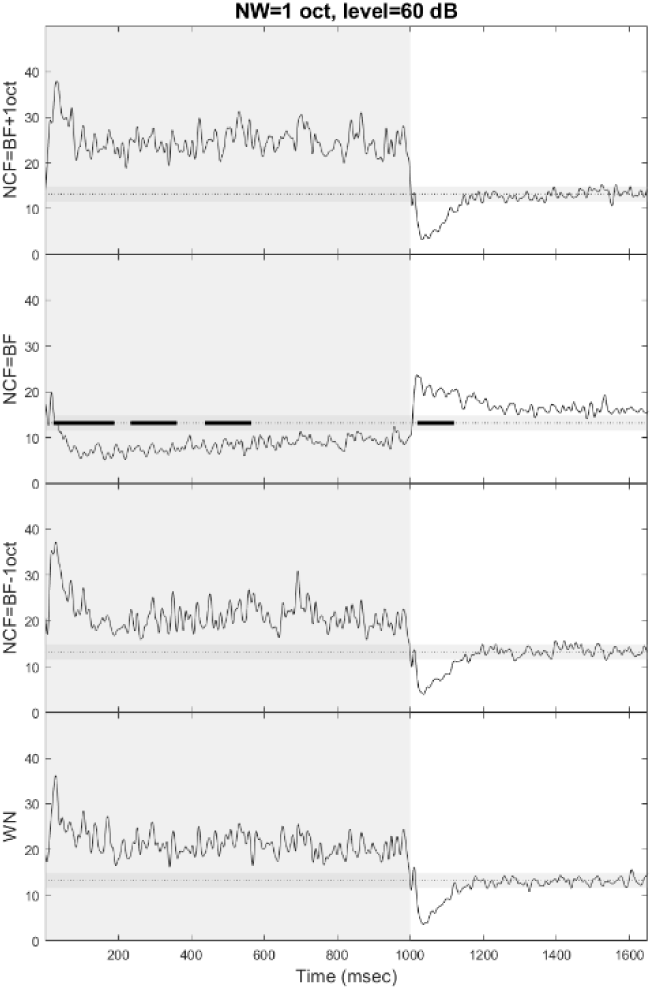
Averaged firing rate during and after NN (notch width (NW) is 1 oct) and WN stimulation (stimulation level is 60 dB). Notch center frequency is expressed as octave relative to cells’ BF (on the left side of each figure). Light gray area indicates sound stimulation and darker gray horizontal band represents the mean +/- 2 * standard deviation calculated from the time period 400-650 msec after WN presentation. Black horizontal bars indicate significant clusters (p&0.05, significant difference between NN and WN, see methods).

Figure 12 shows the averaged neural activity for WN and NN_BF presented at three different levels (40, 60 and 80 dB SPL, notch width was 1 oct) collected from 8 cells. There was no significant differences between the two stimulus conditions at 40 dB. At 60 and 80 dB, the neural activity during NN stimulation was significantly reduced compared to WN (p<0.05). The neural activity was significantly enhanced immediately after the NN_BF presentation at 60 dB only (p<0.05).

**Figure 12.**
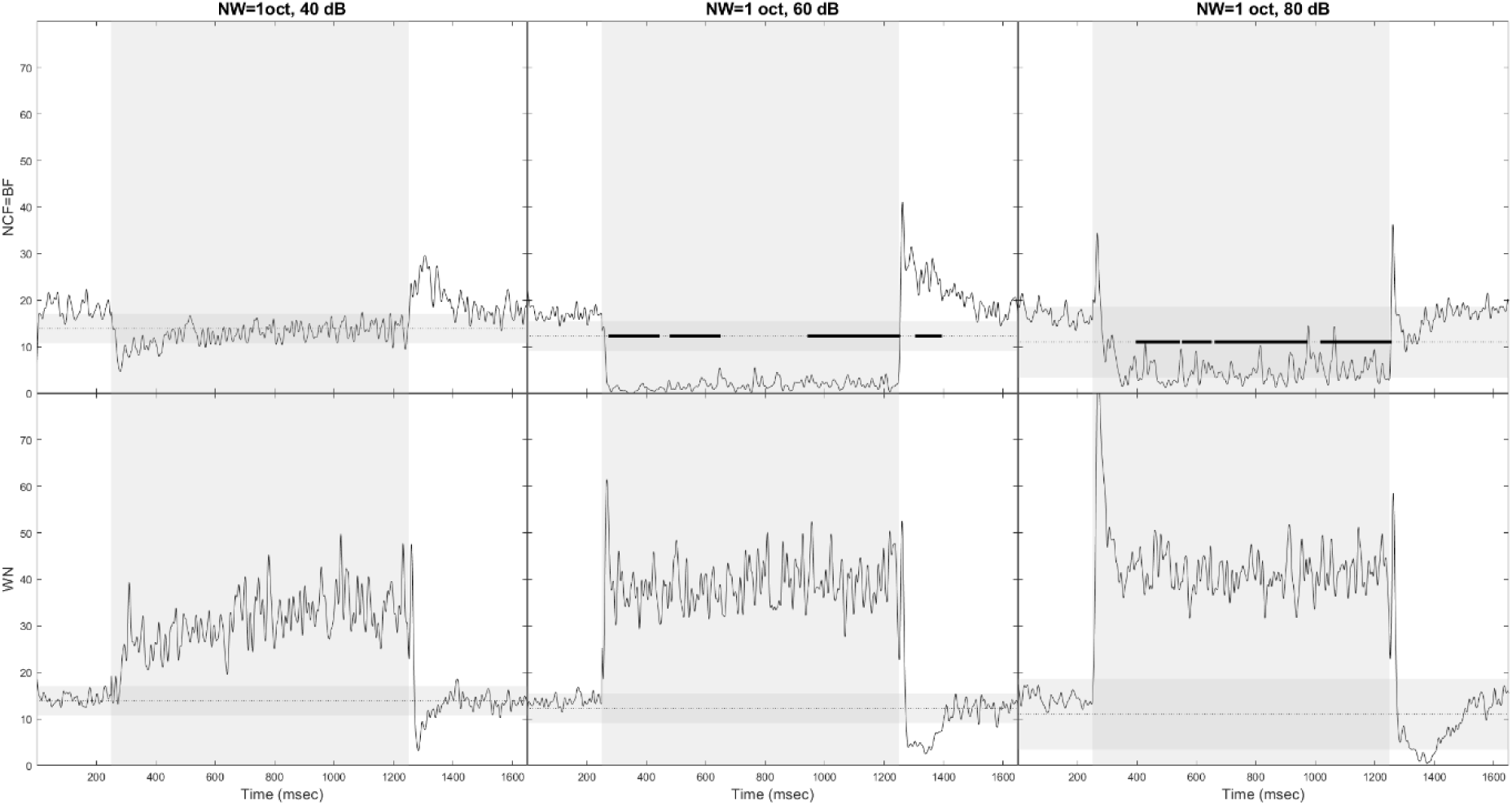
Averaged firing rate during and after NN (notch width is 1 oct and notch center frequency corresponds to BF) and WN stimulation. The sound level is indicated on the top of each column. Light gray area indicates sound stimulation and darker gray horizontal band represents the mean +/- 2 * standard deviation calculated from the time period 400-650 msec after WN presentation. Black horizontal bars indicate significant clusters (p&0.05, significant difference between NN and WN, see methods).

Figure 13 shows the averaged neural activity for WN and 3 conditions of NN_BF (notch width of 0.5, 1 and 2 octaves) collected from 12 cells. Due to the variability across cells and the small number of cells for which the effects of notch width have been documented, the nonparametric statistical test does not show any difference between the NN and the WN. We note, however, the same trend observed in Figure 10 and 11: neural activity is reduced during NN stimulation and is enhanced immediately after NN presentation. Moreover, offset responses tend be larger (in terms of duration) for notch width of 1 oct compared to notch width of 2 oct.

**Figure 13.**
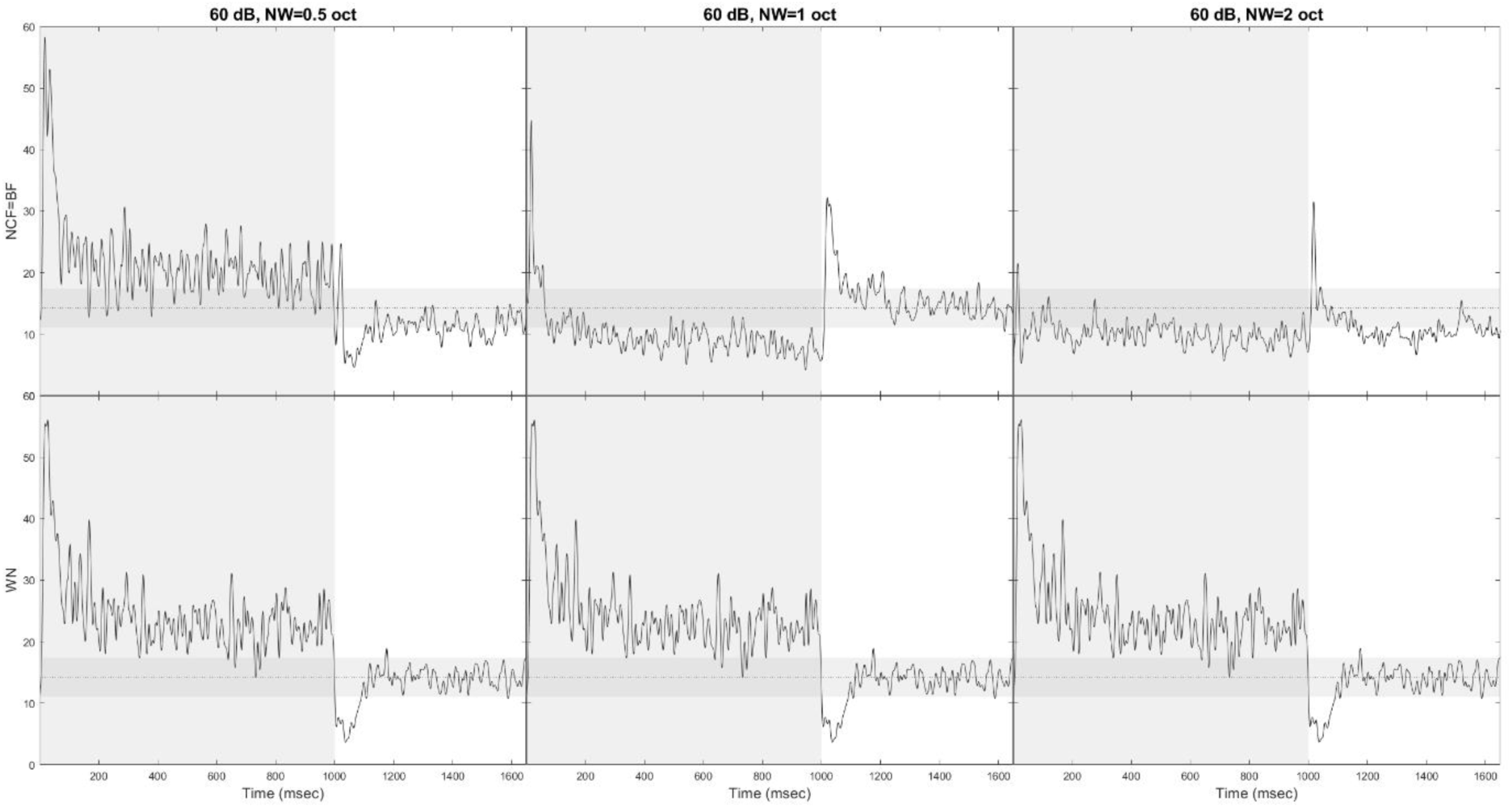
Averaged firing rate during and after NN (notch center frequency corresponds to BF) and WN stimulation (level is 60 dB SPL). The notch width is indicated on the top of each column. Light gray area indicates sound stimulation and darker gray horizontal band represents the mean +/- 2 * standard deviation calculated from the time period 400-650 msec after WN presentation. There was no significant difference between NN and WN (p>0.05).

## Discussion

### Summary of the results

The main goal of this study was to investigate the neural responses in IC during and after NN presentation in order to better understand the mechanisms of auditory enhancement and ZT. Various characteristics of the NNs were studied, namely intensity level (40, 60, 80 dB SPL), notch center frequency and width of spectral notch (0.5, 1 and 2 octaves). Offset responses were much more prevalent after NN stimulation (n=55/99) than after WN presentation (n=16/99). Onset and offset responses were both present on a same recording in only 18 cells.

The present study shows that NN with notch center frequency corresponding to neuron’s BF tends to be associated to a decrease, or even a complete suppression, of neural activity during NN presentation, and a strong offset response after the NN presentation. These effects were dependent on the characteristics of NNs: suppression of neural activity during NN presentation and size of offset responses after NN presentation tend to be maximal at 60 dB SPL and for notch width of 1 octave. Moreover, various types of offset responses could be observed: they could be sharp and/or more spread-out (up to 200-300 msec msec after stimulus presentation). The latency of offset responses after NN presentation is variable: neural activity immediately after NN stimulation can be briefly suppressed after stimulus presentation. The averaged neural activity during WN stimulation is usually increased (sustained activity), while it is temporarily decreased (or even suppressed) up to 200 msec after WN presentation.

### Comparison between our study and the literature

The percentage of cells with offset responses observed in the present study is around 60%, which is slightly below the percentage reported in another study in mice were communication calls were used (Akimov et al., 2017), and well above what has been found in other studies after tone pip presentation (Kasai et al., 2012; Kopp-Scheinpflug et al., 2018; Young and Brownell, 1976). In fact, it is difficult to compare the prevalence of offset responses between studies in a given auditory structure since the presence or absence of offset responses depends heavily on the spectro-temporal characteristics of the stimuli used (Kopp-Scheinpflug et al., 2018; this study). For instance, offset responses depend on whether the stimulus is a tone pip or noise, on the stimulus bandwidth (for noises) and stimulus duration and intensity level (Kasai et al., 2012; Solyga and Barkat, n.d.; Sołyga and Barkat, 2019; Young and Brownell, 1976). Moreover, offset (but also onset) responses are facilitated when the frequency components of communication calls are “resolved” by the mouse auditory system, i.e. each frequency component (first harmonics or formants) falls within an auditory filter (Akimov et al., 2017). Our study adds to this literature on “combination-sensitive” neurons (i.e. neurons responding with maximal strength to specific spectro-temporal features) (Wenstrup et al., 2012) reporting in awake mice that offset responses in IC depends on the presence of spectral notch in a broadband noise, especially when the notch center frequency corresponds to the neuron’s BF.

The pattern of responses observed in our study is reminiscent of the type IV neurons that have been described in the dorsal cochlear nucleus. These neurons have a complex tuning, dominated by large inhibitory regions, with relatively small patches of excitation (Young and Brownell, 1976). In average, they have large spontaneous activity (at around 50 spikes/sec) and can have offset responses (Shofner and Young, 1985; Voigt and Young, 1990). The response map (neural excitation and inhibition as a function of frequency and level of tone pips) does not predict the responses to NNs, emphasizing the non-linear processing of stimulus spectro-temporal characteristics (Spirou and Young, 1991). Type IV neurons receive inhibitory inputs from Type II/III neurons which have inhibitory side bands (Voigt and Young, 1990). Type II/III neurons are strongly excited by tone pips and inhibited by broadband stimuli. As a result, type IV neurons are usually strongly inhibited by tone pips and can be excited by broadband noises. Adding to the non-linear behavior of these neurons, which is also relevant for the present study, type IV neurons can be inhibited by broadband noises with spectral notch (Spirou and Young, 1991; Young et al., 1992). Type IV neurons are thought to be the principal (or giant) cells projecting to the contralateral IC (Adams and Warr, 1976; Spirou and Young, 1991), making them plausible candidates as the source of the offset responses observed in the IC (see below).

Consistent with the present study, we have shown in primary auditory cortex of awake and anesthetized guinea pigs that offset responses are much more prevalent after NN than after WN (Schilling et al., 2023). However, the responses in the IC observed in the present study are very different from what is observed on auditory cortex. First, many neurons show sustained activity during stimulation (not in the auditory cortex), which allows to assess inhibition during stimulation. Second, offset responses in cortex are much shorter and less diverse from what is observed in IC. While offset responses last on average around 200 msec in the IC (according to our study), they rarely last longer than 100 msec in the auditory cortex (Schilling et al., 2023). Therefore, it seems unlikely that the IC is the source of cortical offset responses.

### Mechanisms of offset responses

First, one may wonder whether the responses observed in the IC are inherited from lower levels or whether they are created de novo in the IC. However, extracellular recordings do not allow to directly address this question. As suggested above, the pattern of firing during and after NN stimulation may result from type IV neurons in the dorsal cochlear nucleus. The technique we used in the present study, while it provides clean single cell activity in awake animals, does not allow to record from enough time to document relevant cell properties, including detailed frequency-level tuning properties. Therefore, we cannot tell if the cells we have recorded show all characteristics of type IV pattern. Interestingly, many cells recorded in the present study presented narrow excitatory region over frequency and lateral inhibitory sidebands. The present study was not designed to investigate the putative relationship between the receptive field (obtained from tone pips) and the pattern of responses to WN and NN. However, it seems that the pattern of responses showing neural inhibition during NN stimulation, followed by neural excitation, corresponds to cells with narrow excitatory region and lateral inhibitory side-bands. Consequently, the NN with notch center frequency centered on BF, could recruit the inhibitory sidebands of these cells, while avoiding the excitatory central region. This could hyperpolarize the membrane potential and decrease or suppress neural during stimulation. However, we also recorded from cells that seemed to show non-linear integration of inhibition and excitation (see Figure 2 – fourth column, and supplementary figures). Further analysis is needed to investigate the relationship between frequency-level response maps and the pattern of responses during and after WN/NN presentation.

Considering in more details the underlying mechanisms of the pattern of activity observed in the present study, it is likely that inhibitory processes and post-inhibitory rebound play a critical role. The detailed biophysical mechanisms of post-inhibitory rebound have been addressed in the superior paraolivary nucleus, a structure involved in sound localization (Kopp-Scheinpflug et al., 2018, 2011). In brief, offset responses can emerge from inhibition during stimulation, combined with a large hyperpolarization-activated nonspecific cationic current (I_H_), with a secondary contribution from a T-type calcium conductance (I_TCa_). In our study, most offset responses follow a strong inhibition of neural activity during acoustic stimulation, often below the background activity. It has been shown that post-inhibitory rebound can occur in the IC itself through activation of T-type Ca2+ channels (Sun et al., 2020). T-type Ca2+ channels in IC can be activated after removing its inactivation by membrane hyperpolarization. On the other hand, it has been suggested that h-channels (I_H_) may not be involved in the rebound of activity in IC (Sun and Wu, 2008). However, it has been reported that h-channels intrinsic to IC can play a role in the rebound of neural activity after stimulation (Koch and Grothe, 2003). Interestingly, I_H_ and I_TCa_ contribute differently to the offset responses. Inhibition during stimulation hyperpolarizes the membrane potential, which activates I_H_ and removes I_TCa_ inactivation. While I_H_ play a major role for sharp and short-latency offset responses (see Figures 3 and 10), I_TCa_ is associated to slower depolarization and spiking activity more spread out over time (Kopp-Scheinpflug et al., 2011) (see Figures 2 and -8). One also notes that KCC2 co-transporters play a major role in this cascade of mechanisms involved in offset responses by maintaining low intracellular concentration of chloride. For most of the cells recorded in the present study (see individual examples), I_TCa_ and I_TCa_ likely both contribute to the offset responses, generating short-latency and time-spread offset responses (see Figure 3, where both types of offset responses can be distinguished, for NN_8000Hz). Ultimately, it is the level of neuronal inhibition that triggers the cascade of events leading to neuronal excitation immediately after sound stimulation. A sufficient level of inhibition activates cationic conductances which can produce excitatory rebound responses. Conversely, if inhibition does not exceed a certain hyperpolarization threshold, I_h_ and I_TCa_ remain inactive, and inhibition, which is not counterbalanced by these depolarizing currents, may then persist for several hundred milliseconds after stimulation (see Figures 2, 4, 6 and 8). In summary, the level of hyperpolarization “decides” whether the post-stimulation time window is enhanced (after strong inhibition) or reduced (after mild inhibition) (see below for speculations about the function implications of this mechanism).

Finally, our results do not corroborate the hypothesis proposed to account for auditory enhancement suggesting that adaptation of lateral inhibition plays a major role (Nelson and Young, 2010; Viemeister and Bacon, 1982; Wang and Oxenham, 2016). This hypothesis suggests that neural inhibition, including lateral inhibition directed towards the spectral gap, is adapted (reduced) by the presentation of the masker without the target. Hence, when the target and the masker are presented simultaneously after the presentation of the masker alone, the target frequency undergoes less inhibition and is therefore enhanced compared to the same stimulus not preceded by the masker alone. In the present study, however, we observed that inhibitory processes within the spectral notch are very strong and do not adapt over time. Furthermore, auditory enhancement after stimulus presentation (offset responses in our study) tends to be proportional to the strength of inhibition within the spectral notch, which is contrary to what is predicted by the model based on adaptation of inhibition. In summary, our results suggest that the neural enhancement following notched stimuli is caused by the balance between excitation and inhibition and biophysical mechanisms involving h- and T-type Ca2+ channels that show fundamental non-linear properties: they may be activated after membrane hyperpolarization reaches a given threshold.

### Functional implications

The results presented in this study are very clear and robust, observed in many cells, and may reflect a general mechanism of the auditory system that plays an important functional role. Interestingly, the neurons in IC (like neurons in dorsal cochlear nucleus and auditory cortex) are extremely sensitive to spectral gap in broadband noise. One may wonder why central auditory neurons exhibit this sensitivity to notched stimuli. In other words, what is the significant ecological value of spectral notches that warrants special treatment by the central auditory system? In this context, it is well-known that the spectrum of acoustic stimuli, shaped by direction-dependence of sound propagation through the external ear, provide relevant cues for sound localization. Broadband sounds are filtered by the external ear, which “adds” spectral notches to the sound spectrum. These spectral cues can be used to estimate sound source direction (Anbuhl et al., 2017; Musicant et al., 1990; Rice et al., 1992; Yu and Young, 2000). The present study shows that the activity of neurons with BF corresponding to the notch are reduced. Therefore, a simple neural representation of these monaural cues could simply correspond to the overall activity on the tonotopic axis: relatively low activity in a frequency band surrounded by some background activity in adjacent frequency bands could provide a cue of sound direction. However, this mechanism may come at a price: the frequency band of the spectral notch are neglected while it may be important. In this context, the auditory enhancement after NN presentation can be seen as a sort of “wake-up call” for the neurons encoding frequencies that was initially falling within the spectral notch and inhibited during NN presentation (Sun and Wu, 2008), namely after the acoustic source has moved or after the head or pinna has moved. One can further speculate that animals may use a strategy consisting of “scanning” the acoustic environment by moving the head or pinna, cutting the flow of acoustic inputs into short time windows alternating auditory suppression and enhancement of specific spectral regions.

Offset responses result from mechanisms and neural pathways different from those generating onset responses (Kopp-Scheinpflug et al., 2018; Schilling et al., 2023). It has been suggested that offset responses may play a role in estimating stimulus duration (through the delay between onset and offset responses) (Kopp-Scheinpflug et al., 2018). On our opinion, it is unclear exactly how duration is estimated when the offset responses are so sensitive to the energy and spectral characteristics of acoustic stimuli. Instead, we speculate that offset responses may reflect a collateral effect of a key neural mechanism (“wake-up call” and auditory enhancement) essential for processing rapidly changing acoustic environment, including spectral characteristics added by the head-related transfer function. This hypothesis suggests that post-inhibitory rebound aims to reinforce excitatory inputs (due to new or rapidly changing acoustic stimulation) arriving shortly after a period of synaptic inhibition and that offset spiking activity may not be particularly useful, at least in non-echolocating mammals.

It should be noted that our results could also provide an explanation for the differences in auditory filters shapes obtained from simultaneous and forward masking using the NN method (Leschke et al., 2022; Moore et al., 1987; Oxenham and Shera, 2003). Moreover, the mechanism suggested in our study could also contribute to refine the central representation of frequencies falling in the notch. The post-inhibition rebound of activity is supposed to increase the stimulus pattern of excitation mostly around the center frequency of the notch, while the edges of the pattern of excitation may be unchanged or even reduced by the “post-activation hollow” of neural activity on either side of the target frequency. Consequently, the pattern of excitation should be sharper for targets with frequency close to the notch center frequency.

On another note, it is plausible that the large and spread out offset responses reported in the present study after NN presentation may be a neural correlate of ZT: the brief and faint tonal sensation experienced by human listeners immediately after NN presentation (Franosch et al., 2003; Norena et al., 2000; Wiegrebe et al., 1996; Zwicker, 1964). The characteristic of offset responses reported in the present study are consistent with the properties of the ZT. First, offset responses are much more prevalent and larger when neuron’s BF correspond to the notch center frequency. This is consistent with the ZT pitch falling within the spectral notch. Second, the offset responses described in the present study are relatively long lasting (around 200-300 msec), which is compatible with the ZT duration estimated in human subjects. Finally, offset responses seem to be maximum at 60 dB SPL and for notch width of 1 octave, which is also consistent with what we know from human studies on ZT (Zwicker, 1964, and unpublished observations). We speculate that there is no adaptive value in hearing a ZT after a NN: the ZT, like the offset responses from which it originates, may be perceptual and neural collateral effects of the post-inhibitory rebound generated by the NN presentation, respectively. Studying whether subjects experience ZT and/or whether perception is changed during and after NN presentation, may be used in subjects with or without hearing loss, with or without tinnitus or hyperacusis, to evaluate certain properties of interest of the central auditory system, in particular the putative post-inhibitory rebound, the contribution of I_H_ and I_TCa_ to this rebound and the KCC2 co-transporters to inhibition during stimulation (Bergeron et al., 2014; Hyde et al., 2011; Kadam and Hegarty, 2024; Parameshwarappa et al., 2024).

## Acknowledgements

This research was supported by grant R01 DC016918 from the National Institute on Deafness and Other Communication Disorders of the U.S. Public Health Service. This work has benefited from a french government grant managed by the Agence Nationale de la Recherche under the France 2030 program, reference ANR-23-IAHU-0003.

---

**Supplementary Figure 1.**
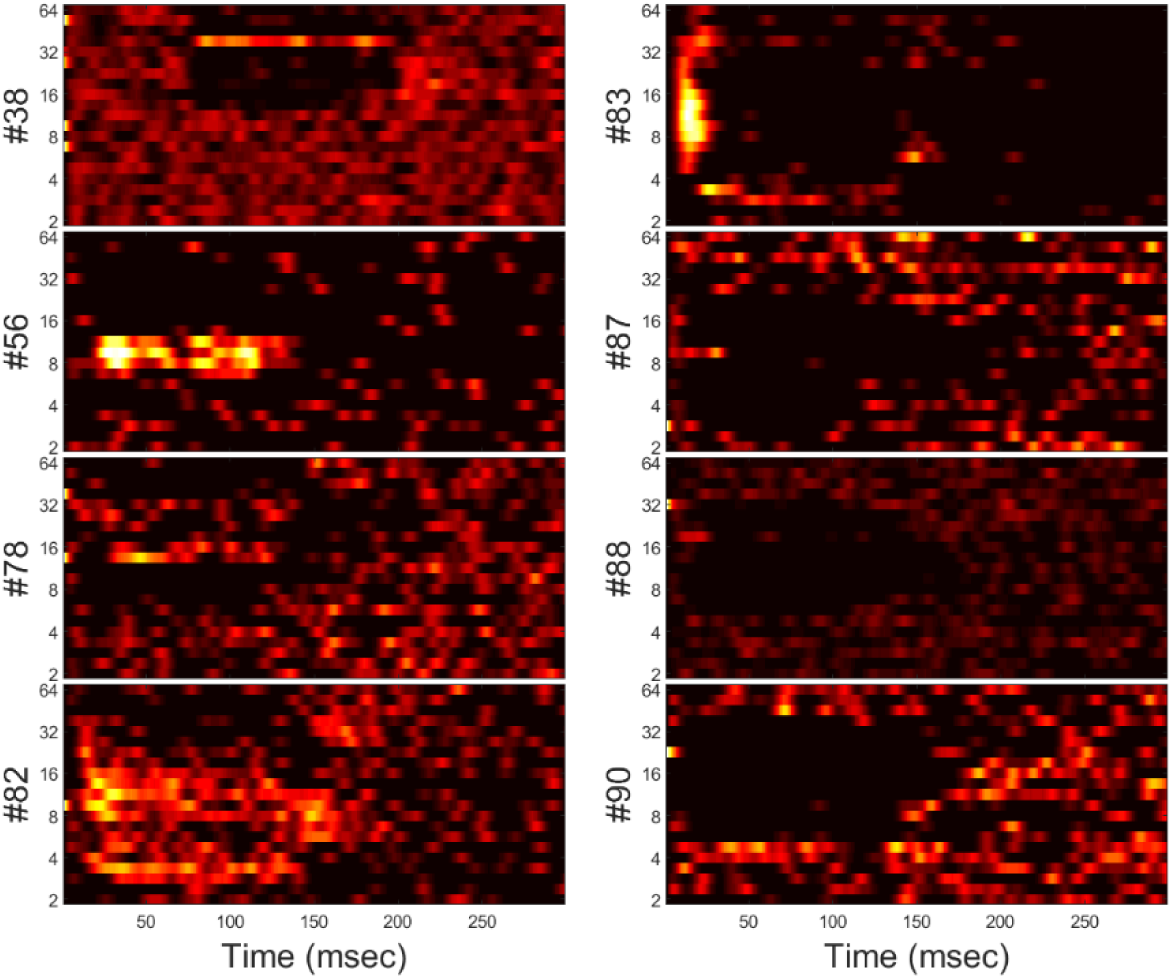
Frequency response as a function of time for all cells whose responses to NN and WN are shown in Supplementary Figures 2-9. Tone pips (100-130 msec, frequency from 2kHz to 64kHz, step of ¼ octave) were presented at 60 dB SPL. In a few examples, tone pips were preceded by 50-70 msec silence. Cell numbers are shown on the left side of each frequency response. The most common pattern of responses is narrow excitatory area surrounded by lateral inhibitory sidebands.

**Supplementary Figure 2.**
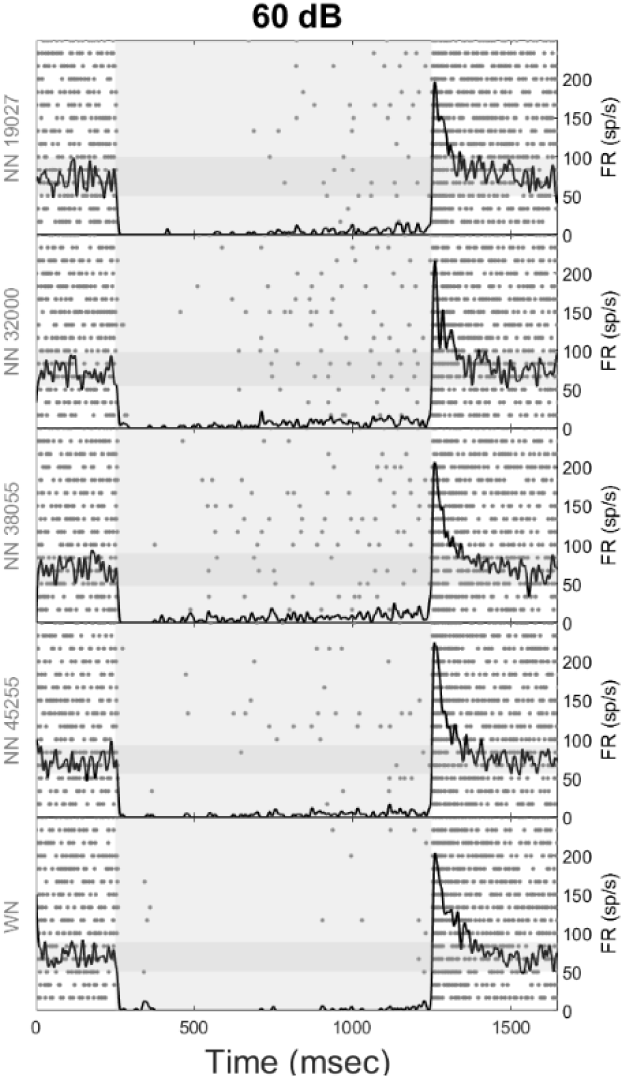
otherwise) is indicated on the left side of each figure. Each gray dot corresponds to an action potential (each row corresponds to a sound sequence, 15 repetitions are shown). The black line corresponds to the firing rate (right y-axis) calculated using a sliding gaussian window of 20 msec. The light gray area corresponds to the stimulus presentation, and the darker gray horizontal band (from 0 to 1650 msec) to the mean spontaneous activity +/- 2 * standard deviation (derived from the time window between 400 and 650 msec after sound stimulation).

**Supplementary Figure 3.**
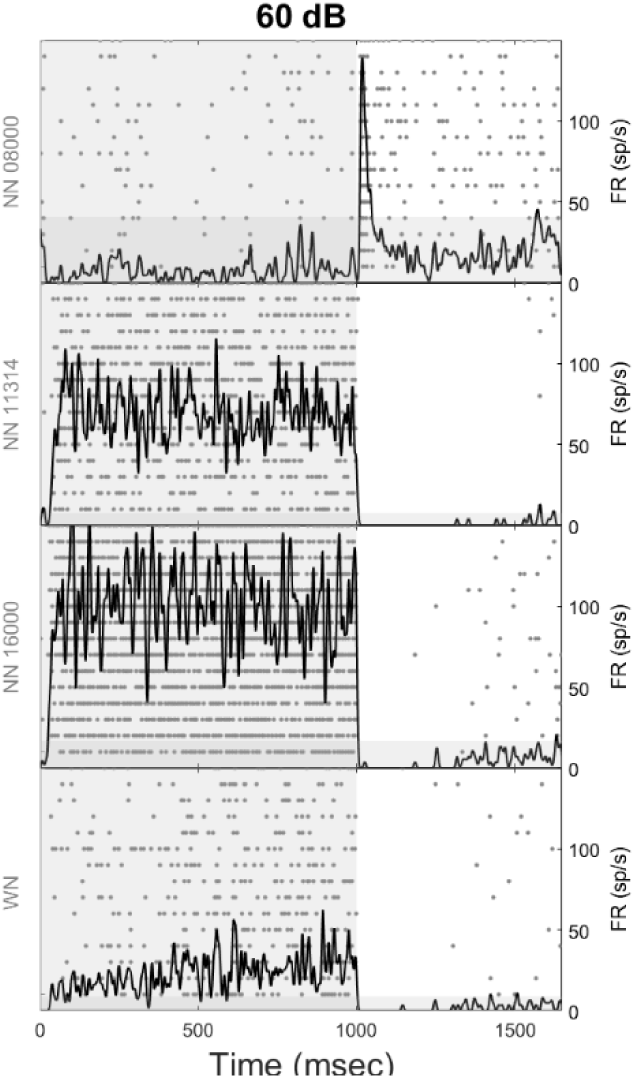
The figure shows the pattern of responses for cell #56. Same than Figure 2 otherwise.

**Supplementary Figure 4.**
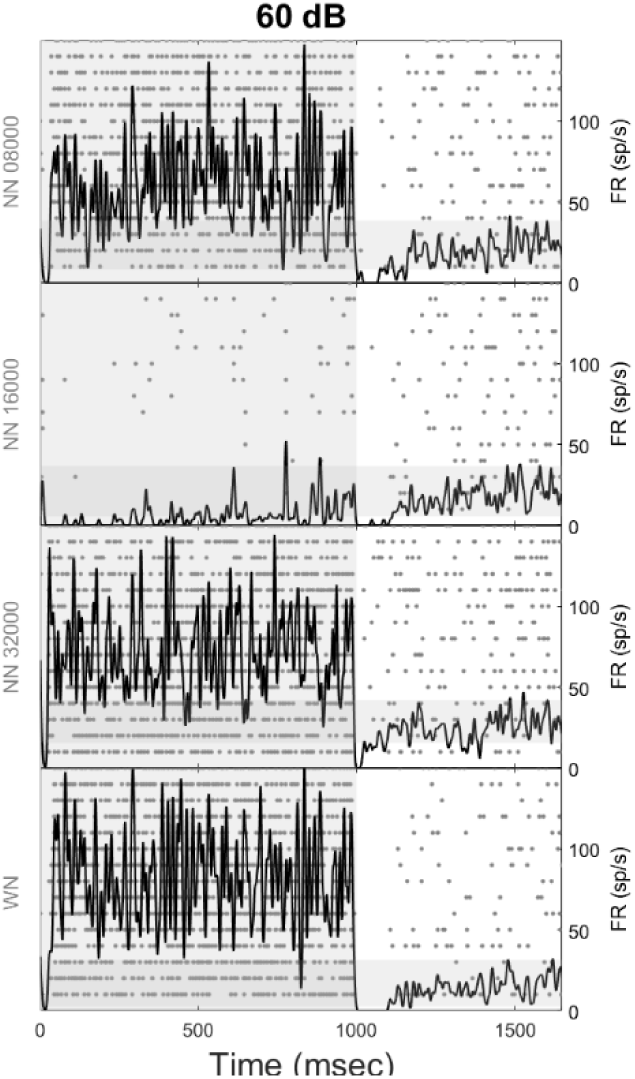
The figure shows the pattern of responses for cell #78. Same than Figure 2 otherwise.

**Supplementary Figure 5.**
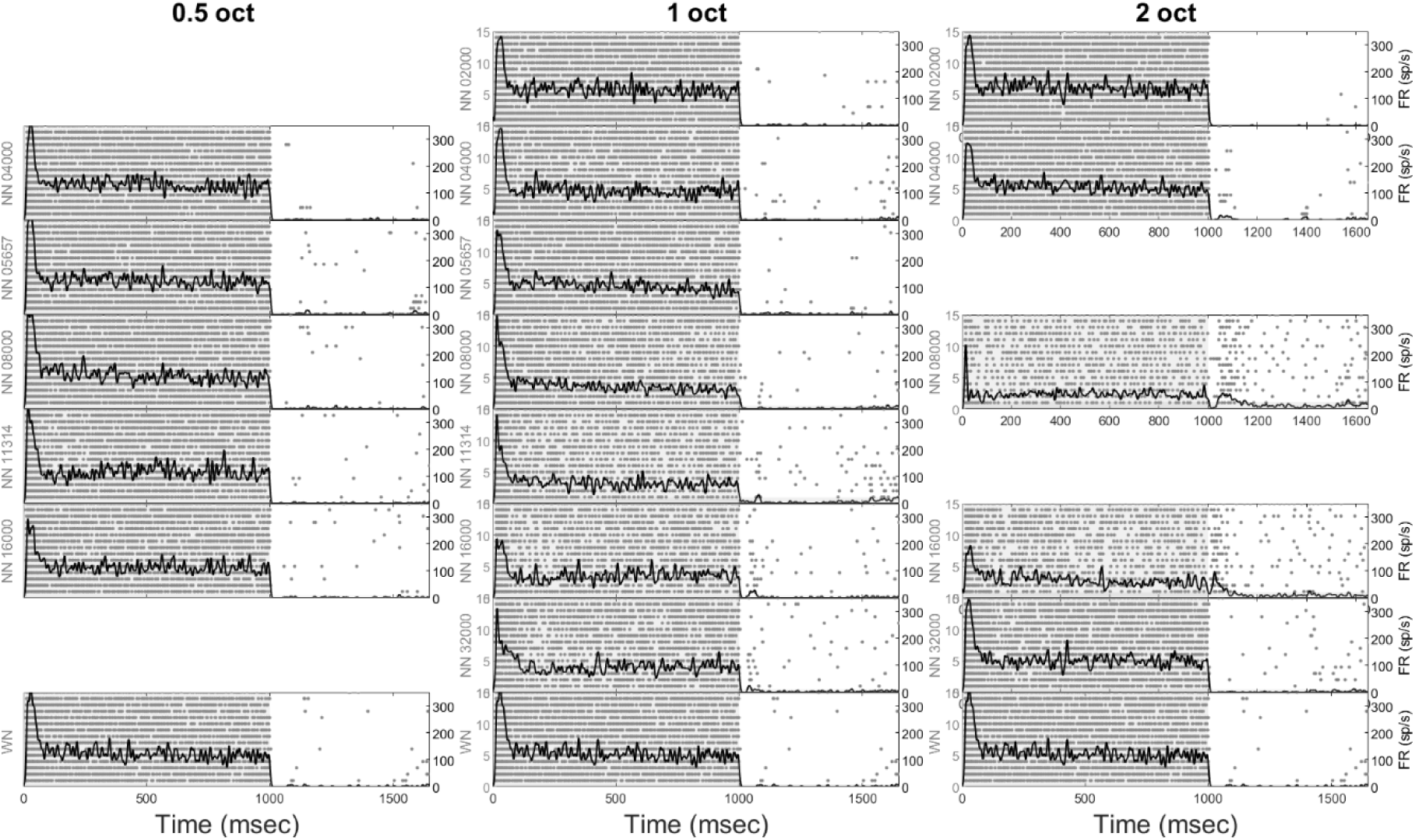
The figure shows the pattern of responses for cell #82. Same than Figure 2 otherwise.

**Supplementary Figure 6.**
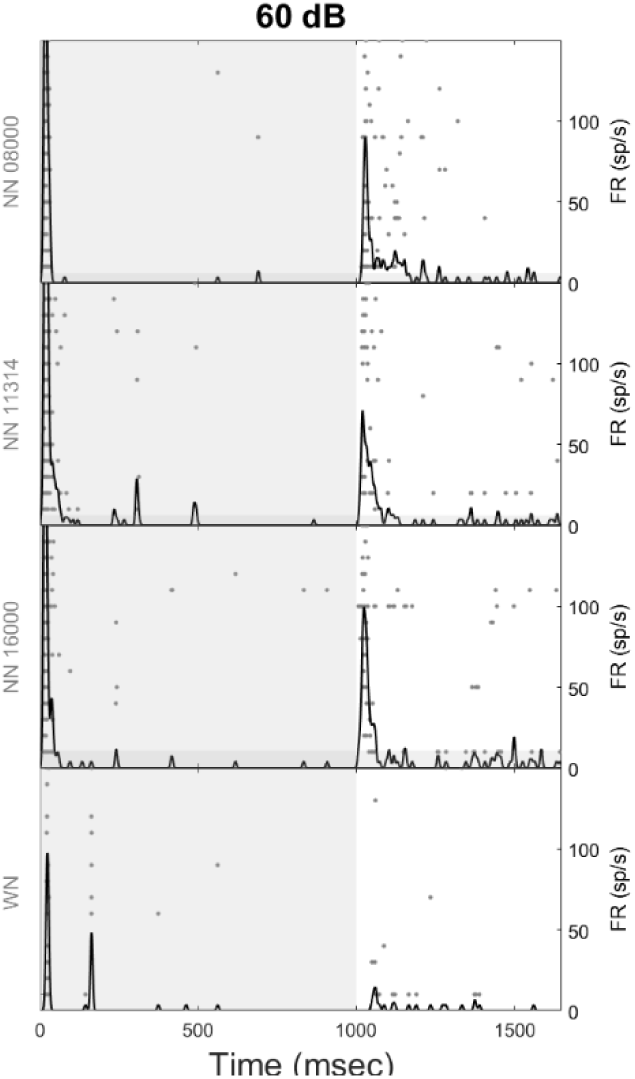
The figure shows the pattern of responses for cell #83. Same than Figure 2 otherwise.

**Supplementary Figure 7.**
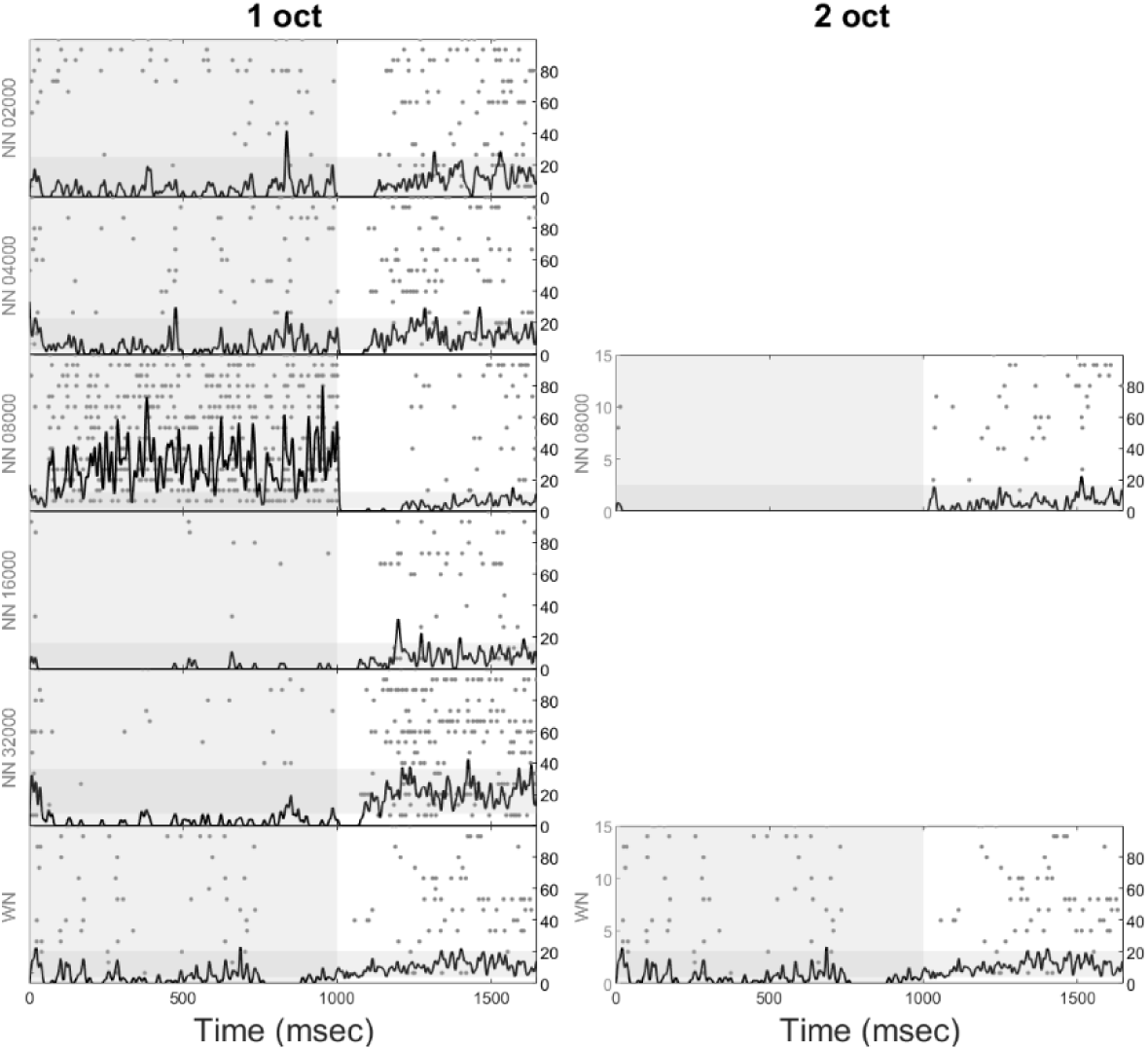
The figure shows the pattern of responses for cell #87. Same than Figure 2 otherwise.

**Supplementary Figure 8.**
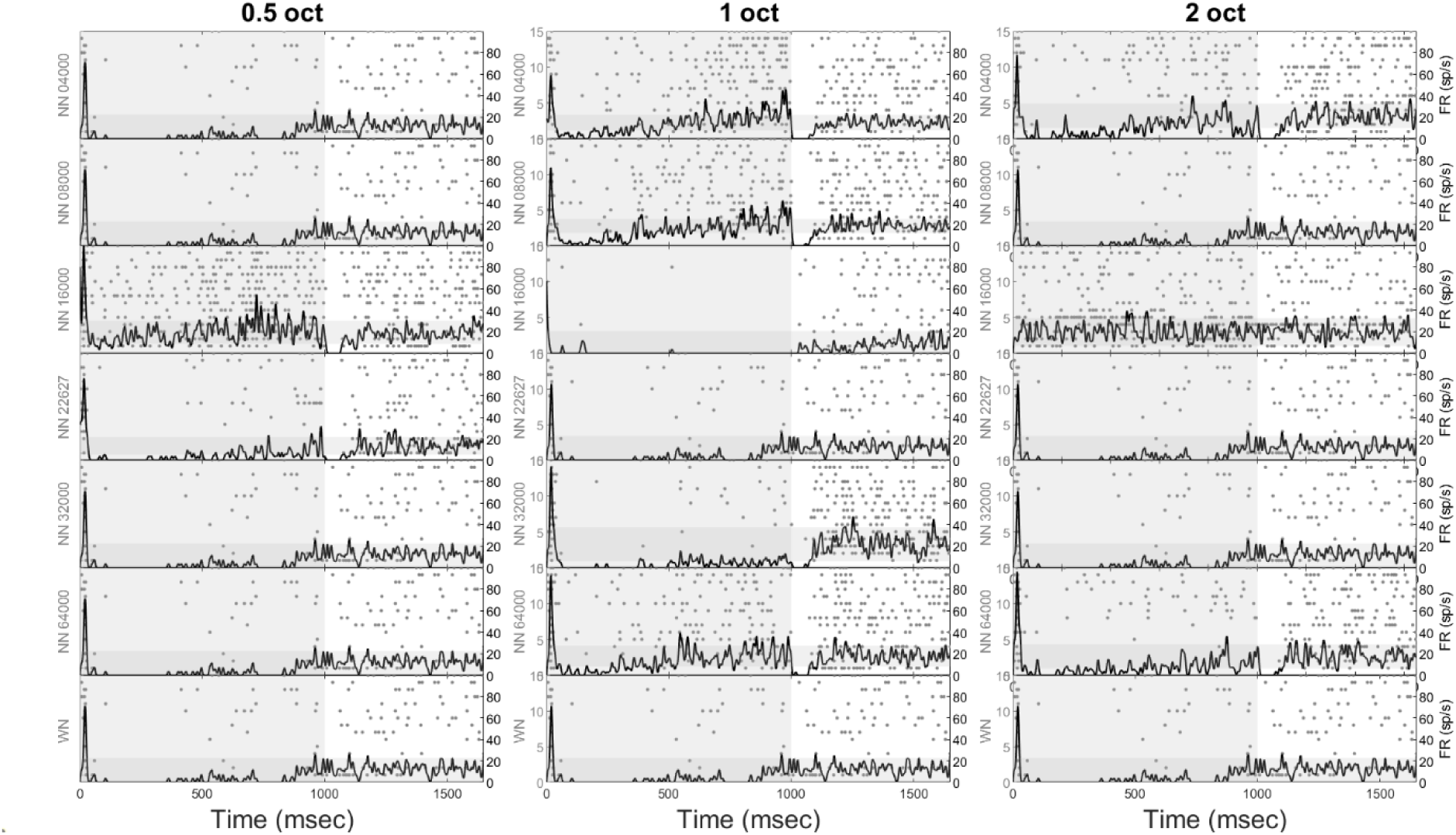
The figure shows the pattern of responses for cell #88. Same than Figure 2 otherwise.

**Supplementary Figure 9.**
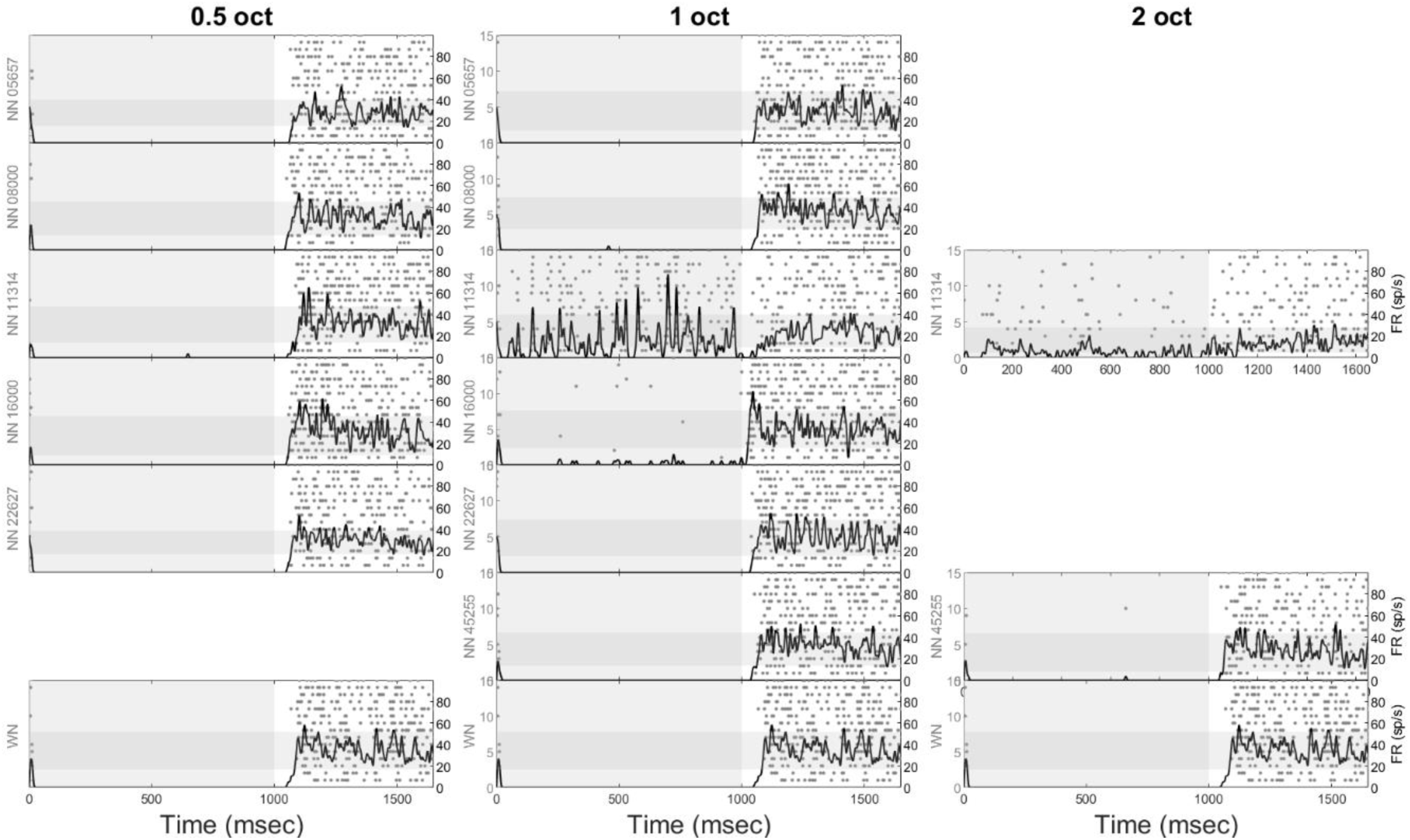
The figure shows the pattern of responses for cell #90. Same than Figure 2 otherwise.

